# Using a Bacterial Protein to Selectively Target Bacterial Biofilms: Treatment of *S. epidermidis* Biofilms with Targeted Photothermal Gold Nanoparticles

**DOI:** 10.1101/2024.09.03.610983

**Authors:** Tanveer Shaikh, Dhanush L. Amarasekara, Radha P. Somarathne, Kenneth Hulugalla, Gabriel J. Alcantara, Madison A. Hejny, Elizabeth R. McCaffrey, Thomas Werfel, Nicholas C. Fitzkee

## Abstract

Biofilm-related infections are associated with high mortality and morbidity combined with increased treatment costs. Traditional antibiotics are becoming less effective due to the emergence of drug-resistant bacterial strains. The need to treat biofilms on medical implants is particularly acute, and one persistent challenge is selectively directing nanoparticles to the biofilm site. Here, we present a protein-based functionalization strategy that targets the extracellular matrix of biofilms. The protein, derived from the extracellular *Staphylococcus epidermidis* autolysin, directs nanoparticles to *S. epidermidis* cell wall components, which are not expected to be present in mammalian tissues. This functionalization is applied to a gold nanoparticle (AuNP) core, along with elastin-like polypeptides (ELPs), which generate a robust photothermal response. In addition to biofilm targeting, the particles exhibit low protein binding, and the photothermal conversion can be modulated by changing the ELP transition temperature. These functionalized AuNPs strongly interact with biofilms under static and flow conditions but exhibit weak interactions with serum-coated surfaces. Near-infrared laser irradiation resulted in a 10,000-fold improvement in killing efficiency compared to untreated controls (p < 0.0001). The targeting strategy utilized here represents a versatile approach to targeting drug-resistant infections and could be readily expanded to other anti-biofilm nanoparticle platforms.

**Graphical Abstract:** 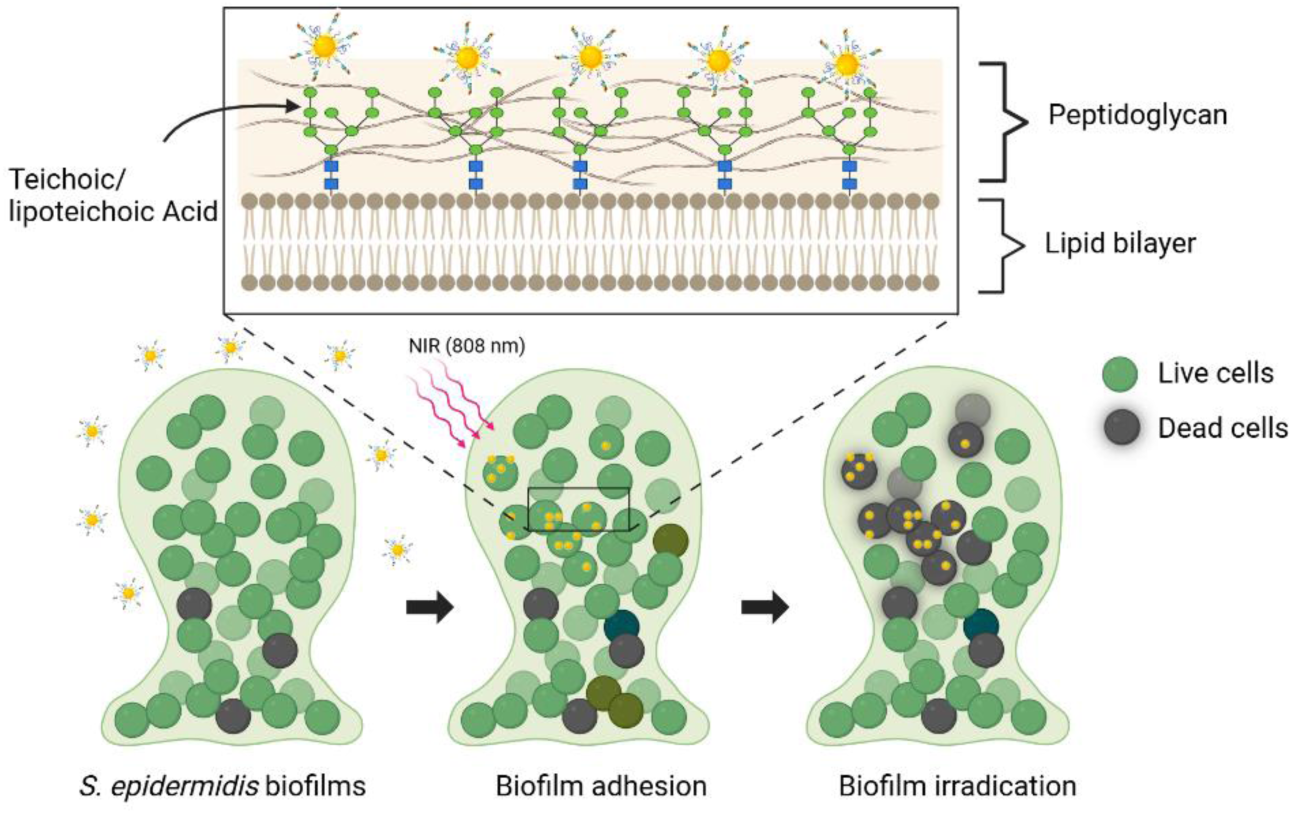

## Introduction

Biofilms ^i^ are structured assemblies of bacterial cells that attach to surfaces and are enclosed in extracellular polymeric substances (EPS).^1–3^ Compared to planktonic bacteria, bacteria in biofilms possess an altered metabolic state and physical environment.^4–6^ *Staphylococcus aureus* and *S. epidermidis* are the primary cause of implant-related biofilm infections, endocarditis, and sepsis.^7^ *S. aureus* produces a wide variety of virulence factors, making it more virulent than *S. epidermidis*.^8^ However, *S. epidermidis* more readily attaches to surfaces and is responsible for 40-45% of infections in hip and knee replacements.^9^ Mera *et al.* reported that the frequency of staphylococcal infections has risen, with hospital stays doubling from 1998-2007.^10^ Therefore, the efficient treatment for biofilm staphylococcal infections is still an immediate and formidable challenge, and developing new therapeutic agents to target staphylococcal biofilms specifically is a prime concern.

While traditional antibiotics are essential in controlling biofilms,^11^ bacterial biofilms can be up to 1,000 times less sensitive to antibiotics than planktonic bacteria.^12^ Drug-resistant biofilms are increasingly common.^10,13^ Nanoparticles hold great potential in resolving the issues of biofilm-targeted drug delivery, and new nanoparticle-based therapies are being developed to complement small-molecule antibiotics. For example, Gao *et al.* prepared azithromycin (AZM) functionalized clustered nanoparticles to treat biofilms. Remarkably, in an acidic biofilm milieu, these clustered nanoparticles can disintegrate and release a secondary AZM-conjugated nanoparticle with a smaller size and positive charge, which helps them penetrate biofilms, boosting antibiofilm activity.^14^ However, the EPS layer functions as a proficient physical and metabolic barrier, resulting in elevated antibiotic resistance by limiting drug penetration and triggering antibiotic inactivation. As a result, several antibiotic-free methods have been developed in recent years, including photodynamic treatment (PDT), photothermal therapy (PTT), and bioactive materials-based antibacterial therapy.^15–18^ Of these methods, photothermal therapy (PTT) has demonstrated growing promise in treating bacterial infections, partly because of the rapid development of photothermal agents (PTAs), the critical component of PTT. In general, PTT uses laser light to produce limited and controlled thermal damage to bacterial cells.^19^ PTT is efficient in treating biofilm infections and sidesteps drug resistance.^20^ However, existing PTAs have a limited ability to target biofilms specifically (e.g., on a medical implant), and the body’s proteins bind to nanoparticles,^21^ forming a protein corona that hinders targeting effectiveness.

Combining multiple functionalization strategies on a single nanoparticle shows promise for developing general-purpose PTAs. For example, Wang *et al*. developed biofilm-responsive caged guanidine nanoparticles (CGNs) to achieve deep biofilm penetration.^22^ CGNs use selective guanidine release, triggered by biofilm acidity, to improve biofilm attachment and penetration. Then, CGNs convert near-infrared (NIR, 800 − 1100 nm) laser energy into localized heat to thermally destroy bacterial biofilms. However, the synthesis of CGNs requires multiple steps, and the guanidino compounds used during the synthesis can induce cytotoxicity and neurological complications.^23^ In another study, Fang *et al*. synthesized α-Fe_2_O_3_/AgAu/polydopamine (PDA) nanospindles, which exhibit antibiofilm activity by combining a controlled Ag^+^ release and a photothermal effect. These particles were 99.9 % effective against *E. coli* and *S. aureus* biofilms.^24^ However, they exhibited poor targeting against biofilms, producing undesirable inflammation in healthy tissues following NIR laser irradiation. Combining multiple functionalities improved PTA performance for both systems, yielding a synergistic benefit.

Previously, we developed thermally responsive nanospheres (TRNs) as a promising platform for treating biofilms under static conditions.^25^ Elastin-like polypeptide (ELP) was used to functionalize citrate-capped spherical gold nanoparticles (AuNPs). This resulted in high photothermal conversion efficiencies (η ≈ 60%) rivaling the values observed for nanorods.^26^ We successfully eliminated both *E. coli* and *S. epidermidis* biofilms under static conditions using TRNs. By tuning the ELP transition temperature (T_t_) we could control the degree to which the temperature increased upon exposure to NIR. TRNs are also biocompatible, employing a citrate-based synthesis method. Nevertheless, the biofilms and NIR treatment were applied under static conditions, and no specific biofilm targeting was employed. A practical system will require a mechanism for targeting biofilms, and it should ideally be effective under dynamic flow conditions, as found in circulating blood.^27^

Here, we aim to address this challenge. Biofilm-targeting TRNs were designed to treat *S. epidermidis* infections (**Scheme 1**), extending the application range of TRNs. Importantly, we use a novel protein-based localization strategy whereby the *S. epidermidis* R2ab domain drives TRN binding to *S. epidermidis* biofilms. R2ab is a subdomain of the autolysin enzyme (AtlE), and it binds to the cell wall, particularly to the lipoteichoic acid (LTA) and wall teichoic acid (WTA) at the septum of dividing cells.^28,29^ R2ab can favorably interact with LTA and peptidoglycan (PGN) in the staphylococcal cell wall, and its normal function supports bacterial cleavage.^28^ Here, we exploit this property so that TRNs can specifically target *S. epidermidis* biofilms. Mammalian cells lack a cell wall, making this an effective strategy. In addition, TRNs developed in this work show low protein binding and high colloidal stability in the presence of serum. We test these targeted TRNs under various in vitro conditions, including those where the nanoparticles are suspended in a flowing solution. We find that the anti-biofilm properties of TRNs are retained even when the TRNs are flowing past established biofilms in a tubular flow cell reactor. Moreover, binding to serum-coated surfaces is relatively low, demonstrating that R2ab-functionalized TRNs have reasonable targeting specificity. The proof-of-concept biofilm targeting TRNs developed in this work could potentially lead to a new class of therapeutic nanoparticles for treating biofilm-associated infections.

## Materials and Methods

### Preparation of 15 nm gold nanoparticles (AuNPs)

Gold nanoparticles were synthesized as described previously.^25,30–32^ The concentrated samples were characterized for size and conformity.^33^ The size of the synthesized AuNPs was measured using dynamic light scattering (DLS; Anton Paar Litesizer 500) and transmission electron microscopy (TEM; JEOL 2100), and they consistently measured 15 nm (Supplementary Material, **Figures S1** and **S2**). The UV–vis spectrum of synthesized AuNPs showed an intense band at 520 nm (Supplementary Material, **Figure S3**), which was used to estimate the AuNPs concentration.^25,33^ Inductively coupled plasma-mass spectrometry (ICP-MS, PerkinElmer ELAN DRC II) was used to determine the Au concentration.^34,35^ The concentrated AuNPs were stored at 4 °C until used. Throughout the text, concentrations are reported using particle concentration (nM) and total Au concentration (ppm). Particles per mL, another common measure of particle concentration, can be calculated from nM by multiplying by 6.0 × 10^11^.

### Preparation of the Elastin-like polypeptide (ELP) construct

The 40-repeat ELP sequence consists of 40 VPGXG repeats, where X is V for all but six repeats. For conjugation to AuNPs, cysteine residues were introduced at position 4 of the first repeat (A4C). The protein was expressed and purified according to previously published methods.^36–38^ The complete protein sequence is available in Amarasekara *et al.*^25^ The protein was stored in 20 mM HEPES, 5 mM NaCl, and 5 mM TCEP at -4 °C for short periods (< 1 month) and lyophilized for longer periods after dialyzing in deionized water.

### Preparation of the GB3 null-fusion (GB3NF) protein construct

The GB3 null-fusion construct (GB3NF) is a small globular protein consisting of the third IgG-binding domain of streptococcal protein G.^39^ It consists of 56 amino acid residues, and a cysteine was introduced in the first β-turn (N8C). The protein was expressed and purified as described previously.^25^ It was stored in 20 mM HEPES, 5 mM NaCl, and 5 mM TCEP.

### Preparation of the R2ab fusion (R2abF) construct

The R2ab fusion protein (R2abF) consists of three regions: a GB3 domain, a linker, and an R2ab domain (Supplementary Material, **Figure S4**). The GB3 domain is similar to GB3NF, containing an N8C variant to facilitate AuNP attachment. The N-terminus of the GB3 domain contains a 6X histidine tag and a thrombin cleavage site to aid purification using affinity chromatography. The linker region was designed to separate the GB3 and R2ab domains, providing a flexible linker and minimizing interaction between R2ab and the (GB3-coated) AuNP surface. The R2ab domain was fused at the C-terminal end of the linker. The fusion protein was subcloned into a pET-15b vector for recombinant protein expression in *Escherichia coli* under the control of the T7 promoter. Plasmid construction was performed by GenScript (Piscataway, NJ). Purification of R2abF was performed using the approach described previously for the isolated R2ab domain.^40^ After purification, R2abF was dialyzed into a buffer containing 20 mM HEPES, 5 mM NaCl, and 5 mM TCEP. The structure of the R2ab and GB3 domains was confirmed by comparing the ^1^H – ^15^N HSQC spectrum of R2abF with the spectra of the isolated domains (Supplementary Material**, Figure S5**). Chemical shift perturbations were minor in the fusion protein.

### Synthesis of biofilm-targeting TRNs (AuNP@PEG5K@R2abF@ELPA4C)

Thiolated polyethylene glycol (PEG) with an average molecular weight of 5,000 was used to coat AuNPs as described previously.^25^ AuNP (AuNP@PEG5K) with different AuNP/ PEG ratios were initially prepared and mixed with R2abF protein to find the optimum conditions to prevent aggregation caused by direct interaction of R2abF with AuNP surface. Here, the R2abF concentration was kept constant (AuNP/ R2abF = 1: 250), and shortly after mixing R2abF with the AuNP@PEG5K, the mixture was vortexed several times to prepare the AuNP@PEG5K@R2abF in 20 mM HEPES at pH 6.5 and 5 mM NaCl. Finally, the desired concentration of ELP was added to the mixture to prepare AuNP@PEG5K@R2abF@ELPA4C. After incubation for 2 h, the solutions were centrifuged three times at 21,300 *g* for 24 min to remove unbound PEG5K, R2abF, and ELPA4C, which remained in the supernatant. The pH of the prepared solutions was confirmed by pH-indicator strips (EMD Millipore) using the supernatant. After the third wash, the pellet was redispersed in 20 mM HEPES pH 6.5 and 100 mM NaCl to provide a suitable environment for phase separation. AuNPs were modified with thiolated PEG5K, GB3NF, and ELPA4C separately to form AuNP@PEG5K, AuNP@GB3NF, and AuNP@ELPA4C as controls. The final concentration of Au was determined using ICP-MS, as described above. DLS was used to assess the nanoparticle hydrodynamic diameter (D_H_), polydispersity index (PDI), and zeta potential. The behavior of prepared AuNP@PEG5K@R2abF@ELPA4C nano-assemblies was characterized using TEM (Supplementary Material, **Figure S7**).

### Interactions of biofilm-targeting TRNs with S. epidermidis biofilms

*S. epidermidis* cells were incubated in brain heart infusion (BHI) medium at 37 °C and allowed to grow for 48 h on a 96-well plate. Following the incubation, the excess planktonic cells were removed by gently washing the plate three times with phosphate-buffered saline (PBS) buffer. Then AuNP@PEG@R2abF@ELPA4C (100 µL) at various concentrations was added to each well and incubated for 12 h at 37 °C. Then, each well was once again washed three times with PBS, and the presence of AuNP@PEG@R2abF@ELPA4C was measured using a Cytation5 plate reader (Biotec) at 25 °C by monitoring the extinction at 520 nm. The morphology of bacterial cells in the presence and absence of AuNP@PEG@R2abF@ELPA4C was observed using scanning electron microscopy (SEM) and energy-dispersive X-ray spectroscopy (EDS; Supplementary Material, **Figure S8** and **S9**) as previously described.^25^ ICP-MS was used to measure the total gold content in each sample well. Each well was treated for 12 hours with aqua regia and then diluted to 10 mL with ultrapure water. Standard curves were generated using several dilutions of a commercial gold standard (Fluka #38168). Samples were analyzed using a PerkinElmer ELAN DRC II ICP-MS system with an autosampler. The total gold concentration in each sample (ppm or μg mL ^-1^) was calculated by interpolating the standard curve.

### Thermoresponsive behavior of bacterial targeting TRNs

The aggregation of the AuNP@PEG@R2abF@ELP at various temperatures was examined using DLS.^25^ Measurements of agglomeration reversibility were performed below and above the transition temperature (T_t_) of AuNP@PEG5K@R2abF@ELPA4C system after equilibrating for 15 min for 5 cycles. UV–vis measurements were obtained before and after 5 cycles to see any structural changes to the AuNP@PEG5K@R2abF@ELPA4C variant during heating and cooling cycles (Supplementary Material**, Figure S10**).

### Transmission electron microscopy (TEM)

On Formvar-coated copper grids, 5 µL aliquots of a 2 nM (24 ppm of Au) AuNP@PEG5K@R2abF@ELPA4C solution were deposited. The grids were kept at the T_t_ and then exposed to NIR irradiation at 808 nm laser at 1.8 W until the excess liquid evaporated. Prepared grids were imaged using a JEOL 2100 with an accelerating voltage of 200 kV (Supplementary Material, **Figure S12**). TEM was performed at the Institute for Imaging and Analytical Technologies (I2AT) at Mississippi State University.

### Photothermal effect of bacterial targeting nano-assemblies

AuNP@PEG5K@R2abF@ELPA4C samples (400 μL) at different gold concentrations (120 – 360 ppm of Au) in a disposable cuvette were incubated at their T_t_ for 15 min and then exposed to irradiation at 808 nm laser at 1.8 W cm^-2^. This laser power was selected based on the temperature changes observed in our previous work.^25^ The sample temperatures were recorded using an infrared thermographic camera every 100 ms (Optris PI400i). To calculate the photothermal conversion efficiency (ɳ) of AuNP@PEG5K@R2abF@ELPA4C, 1 mL of 20 nM (240 ppm of Au) samples were incubated at their T_t_ for 15 min and then exposed to laser irradiation (808 nm, 1.8 W cm^-2^). When the temperature reached the maximum, the laser was switched off. The parameter ɳ was calculated as previously reported.^25,41–43^

### In vitro antibiofilm activity of bacterial targeting nano-assemblies irradiated by NIR light

*S. epidermidis* was cultured in BHI media at 37 °C and allowed to grow overnight with shaking. A seed culture was prepared from the overnight culture with OD_600_ of 0.2, and 100 μL was added to 96-well plates. Biofilm formation was accomplished by incubating the plate statically at 37 °C for 48 h. Following the incubation period, the extra cells were removed by washing three times with 100 μL of PBS buffer. Following the creation of the biofilm, the wells were treated with 100 μL of nanoparticle or control solutions, maintained at 37 °C for 15 min, and then subjected to 808 nm laser irradiation at 1.8 W cm^-2^ for 5 min. After NIR irradiation, the number of colony-forming units was determined as described below.

### Dynamic biofilm formation on plastic tubes using a 3-D printed flow system

A Crealty3D CR-10 printer (Shenzhen Creality 3D Technology Co., Ltd., Shenzhen, China) was used to 3D print the flow cell reactor. The design of these devices was created using the Autodesk Fusion360 3D CAD Software (Online version). Design files for the flow cell are available as part of the datasets stored online (see Zenodo link below). Before use, all components of the flow reactor were sterilized by immersion in ultrapure water and autoclaving for 15 min using the wet cycle. The system was built to accommodate three sterile, 3 cm untreated polystyrene tubes in series (cut from a KIMBLE serological pipette). BHI medium was inoculated with *S. epidermidis* strain 1301.^44^ A 12 V peristaltic pump (ALAMSCN #AL12537) circulated inoculated medium through the system for 48 h at 37 °C, establishing a biofilm on the plastic tubes. After each experiment, the system was washed with sterile PBS for 5 min to remove any excess medium. Visible biofilm formation was observed on the inner walls of the plastic tubes. Biofilm formation on the inner walls of the plastic tubes was further confirmed using crystal violet staining and SEM imaging (Supplementary Material, **Figure S13**). Next, 20 mL of AuNP, AuNP@PEG5K, and AuNP@PEG5K@R2abF@ELPA4C (30 nM or 360 ppm of Au) were circulated through the flow cell reactor separately for 12 h to determine the affinity of these nano-assemblies on *S. epidermidis* biofilm under dynamic flow conditions. The treated tubes with AuNP@PEG5K and AuNP@PEG5K@R2abF@ELPA4C were then subjected to 808 nm laser irradiation at 1.8 W cm^-2^ for 15 min.

To estimate cell viability after treatment, each plastic tube was immersed in 1 mL of PBS medium in a well of a 24-well plate for 30 min with shaking, allowing the live cells to shed into the PBS. Next, the tubes were removed from each well, and the number of colony-forming units was determined using a standard dropwise assay, as described below.

### Determination of colony-forming units per mL (CFU mL^-1^) in static biofilms

A standard dropwise assay was used to determine bacterial cell viability after treatment of static biofilms. Cells were grown in sterile 96-well plates in BHI media, as described above. After NIR irradiation, the nanoparticle solution was removed from each well of a 96-well plate, and 100 μL of sterile PBS was added to each well. Then, biofilms were carefully resuspended in each well by mixing with a pipette. The suspensions were serially tenfold diluted with sterile PBS, and 10 μL of the diluted samples were applied to agar plates. The plates were tipped, allowing the applied droplets to spread along a 2 cm line. After incubation at 37 °C for 18 h, the colonies formed in each droplet were counted. The first serial dilution droplet where 10-50 isolated colonies had formed was used to estimate the CFU mL^-1^ of the initial resuspended biofilm sample.

### Confocal Microscopy Analysis of Biofilm Penetration

Bacterial biofilms were grown on 8-well cell culture slides (CELLTREAT Scientific Products #229168). Two bacteria were used: *S. epidermidis* strain 1301 and a GFP-expressing *S. aureus* strain developed by Cobb *et al*. (ATCC 6538-GFP).^45^ Initially, 1 mL of culture containing 10^8^ CFU mL^-1^ was added to each well, and biofilms were grown statically at 37 °C for 48 hours. Following biofilm formation, the slides were gently washed with sterile PBS to remove non-adherent bacteria. *S. epidermidis* biofilms were stained with 100 µL of 0.1% w/v Hoechst 33258 (Invitrogen #H21491) for 1 hour at room temperature in the dark, then washed three times with 1 mL PBS to remove excess stain. For fluorescent labeling of TRNs, R2abF proteins (2 mg mL^-1^ in PBS, pH 7.4) were conjugated with Texas Red (VWR #76480) using NHS ester chemistry. Briefly, R2abF protein was mixed with Texas Red dye at a 1:10 protein:dye molar ratio in 500 μL of 0.1 M sodium bicarbonate buffer (pH 8.3). The reaction was incubated for 2 hours at room temperature with gentle rotation in the dark. Unreacted dye was removed using a Clarion 10-S25M desalting column (Sorbtech #804003) equilibrated with PBS. Once fluorescently labeled targeted TRNs were prepared, both *S. epidermidis* and *S. aureus* biofilms were treated with the 100 µL of 20 nM fluorescent nanoparticles for 1 hour. After incubation, the biofilms were washed three times with 1 mL PBS to remove unbound and surface-associated nanoparticles. Confocal imaging was performed using an Axiovert 200M inverted microscope (Carl Zeiss) equipped with filter sets for GFP (excitation 488 nm, emission 510 nm), Hoechst 33258 (excitation 350 nm, emission 461 nm), and Texas Red (excitation 595 nm, emission 615 nm). Z-stack images were acquired at 3 μm intervals to analyze biofilm thickness and assess the three-dimensional penetration of nanoparticles within the biofilm structure. Images were processed using Zen 2009 software.

### Cell Viability Assay

The CellTiter-Glo (Promega) luminescence assay was used to determine the cytocompatibility of various AuNP formulations. HEK-293 cells (5 × 10³ per well) were seeded in a 96-well plate with 200 μL of Dulbecco’s Modified Eagle Medium (DMEM) supplemented with 10% fetal bovine serum (FBS) and 1% antibiotic. The plate was incubated overnight at 37 °C, 5% CO₂ to allow for cell attachment. Following incubation, the culture medium was replaced with 200 μL of fresh medium containing different concentrations (10, 20, and 40 nM) of the AuNP formulations, specifically: AuNPs, AuNP@PEG, AuNP@GB3, AuNP@ELPA4C, AuNP@PEG@R2abF, and AuNP@PEG@R2abF@ELPA4C. Cells were exposed to the nanoparticles for 24 hours, after which they were washed three times with PBS to remove excess particles. Post-washing, 100 μL of fresh medium and 100 μL of CellTiter-Glo reagent were added to each well in a 1:1 ratio. The plate was then subjected to 2 minutes of agitation on an orbital shaker to promote cell lysis, followed by a 10-minute incubation at room temperature to stabilize the luminescent signal. The resulting luminescence was measured using a BioTek microplate reader (BioTek, USA). All experiments were conducted in triplicate, and the measured cell viability was compared against untreated control cells.

### Statistical analysis and data availability

Unless otherwise stated, all measurements are reported as the average and the standard error of the mean for at least three independently prepared samples. When technical replicates are used instead, this is explicitly noted in the text and figure legends. Unless otherwise noted, comparisons were performed using one-way ANOVA with GraphPad Prism 10 software. Pairwise post-hoc testing was performed using Tukey’s multiple comparisons test, and statistical significance was reported at the α = 0.05 significance level. All raw data from this study, including 3D printing design files, have been submitted to Zenodo and are available at https://doi.org/10.5281/zenodo.10887304.

## Results and Discussion

**Scheme 1.**
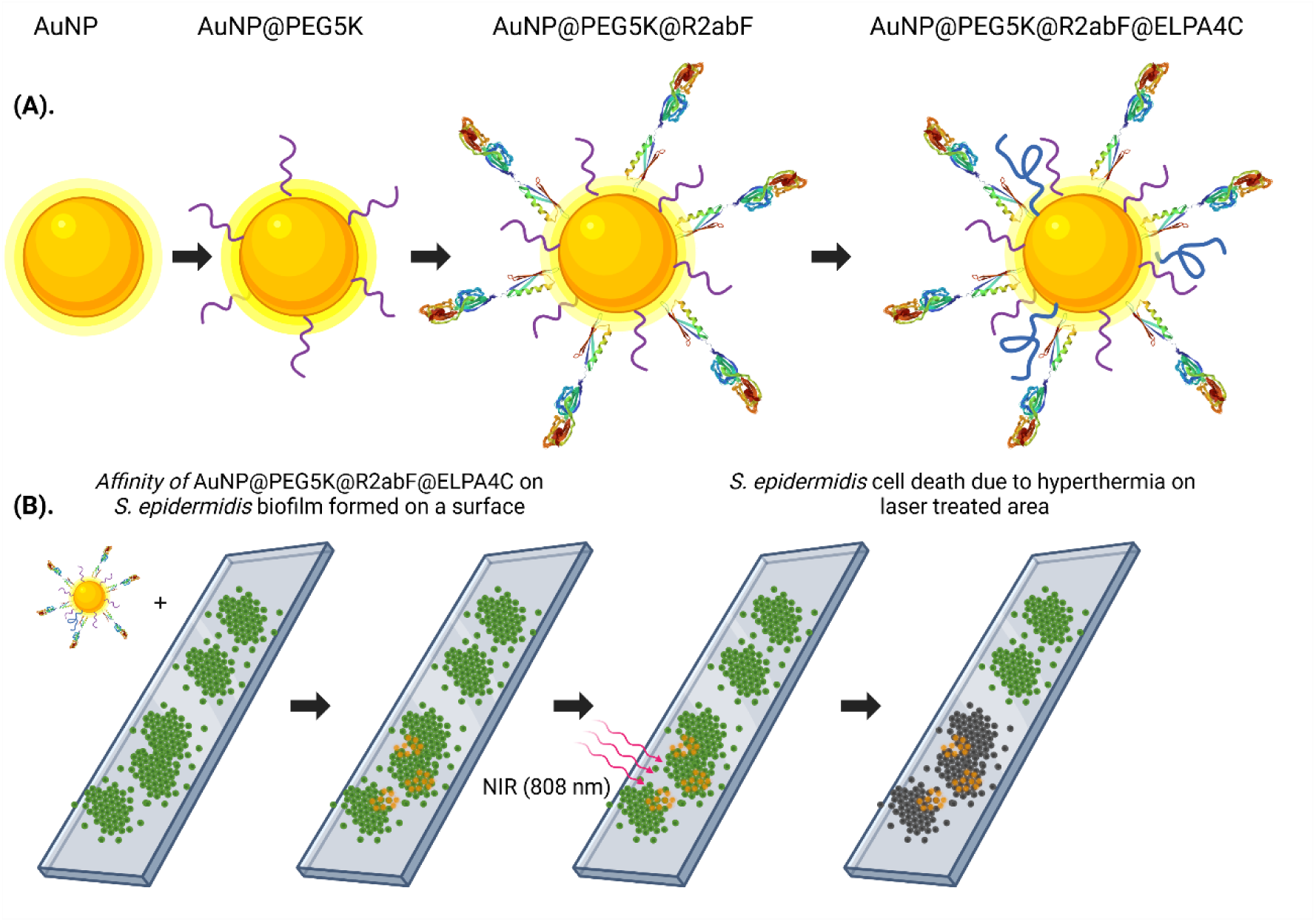
(A) Schematic illustration of the stepwise synthesis of the thermally-responsive nanosphere (TRN) assembly AuNP@PEG5K@R2abF@ELPA4C, (B) Application of final nano-assembly AuNP@PEG5K@R2abF@ELPA4C for targeting and killing *S. epidermidis* biofilm using 808 nm near-infrared (NIR) laser light.

### Synthesis and characterization of biofilm-targeted TRNs

We report our stepwise synthesis of bacteria-targeting, thermally responsive nanosphere (TRN) in **Scheme 1A**. For the synthesis, 15 nm diameter spherical citrate-AuNPs were used (Supplementary Material**, Figure S1** and **S2**). R2ab was chosen to functionalize the AuNPs (AuNP@R2ab), enabling AuNPs to bind the cell wall components of *S. epidermidis*. However, a major challenge with these functionalizing techniques is the potential of losing protein activity upon interaction with the surface of the nanoparticle.^46–48^ For this reason, we used a protein fusion strategy to optimize the orientation and accessibility of the LTA/WTA binding site when conjugating R2ab to the nanoparticle surface.^49^ A novel R2ab fusion (R2abF) construct was engineered that combined R2ab’s biofilm-targeting function with another protein that binds tightly to AuNPs via a strong gold-thiol (Ag-S) linkage. For the AuNP-binding protein, we used a N8C variant of GB3 (a globular protein with 56 amino acid residues), which is known to bind citrate-AuNPs.^30,50–52^ A linker was used to separate R2ab from GB3, reducing the degree to which R2ab interacts with the AuNP surface.^30^

Our initial attempts to create a biofilm-targeting AuNP were unsuccessful. Initially, we treated 15 nm AuNPs directly with R2abF (AuNP@R2abF). This particle rapidly aggregated as determined by a large hydrodynamic diameter (D_H_; **Figure 1A**, gray bars). We hypothesized that the R2ab was binding strongly to the AuNP surface and unfolding, leading to reduced colloidal stability. To control this behavior, we used thiolated polyethylene glycol (PEG) with a 5,000 average molecular weight (PEG5K) to PEGylate the AuNP surface, which increased colloidal stability (**Figure 1A**, remaining bars). The experimentally measured PEG density on these AuNP-PEG conjugates was 0.155 ± 0.005 PEG/nm^2^ (**Figure S6**). These values are lower than those of the prior studies where AuNPs were functionalized exclusively with PEG-SH,^30,53^ suggesting that both R2abF and PEG5K are coating the AuNP surface. The polydispersity index (PDI%) and D_H_ were measured for multiple ratios of AuNP, PEG, and R2ab to determine the optimum formulation of AuNP@PEG5K@R2abF. As the AuNP: PEG5K ratio was increased beyond 1:125 (**Figure 1A**, red bar), no significant reduction in the D_H_ was observed, and the PDI% remained constant; therefore, a AuNP: PEG5K:R2abF ratio of 1:125:250 was chosen as the optimum.

**Figure 1.**
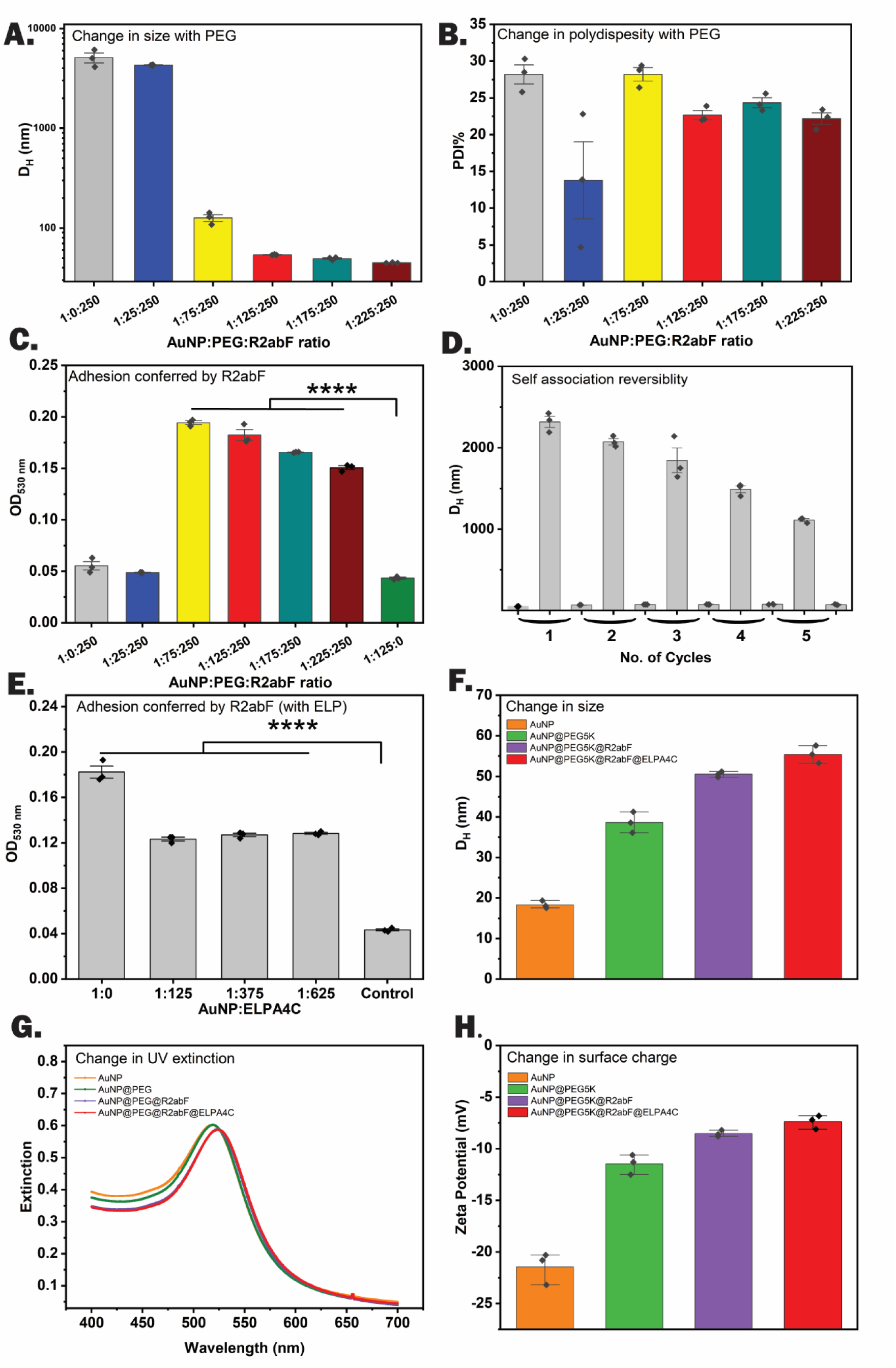
Physiochemical characterization of AuNP@PEG5K@R2abF@ELPA4C. (A.) DLS profiles of AuNP@PEG5K@R2abF to optimize the PEG5K ratio. D_H_ denotes hydrodynamic diameter (nm) and (B.) PDI% denotes polydispersity index as a percentage. (C.) Affinity profiles of AuNP@PEG5K@R2abF on polystyrene surfaces measured by plate reader using λ = 530 nm. (D.) DLS measurements on AuNP@PEG5K@R2abF (20 nM, or 240 ppm Au) for 5 cycles of heating/cooling. A reversible, temperature dependent agglomeration is observed. (E.) Effect of different ELPA4C concentrations on affinity profiles of AuNP@PEG5K@R2abF@ELPA4C on polystyrene surfaces measured by plate reader using λ = 530 nm. (F.) Hydrodynamic diameter (D_H_, nm), (G.) UV–vis extinction, and (H.) Zeta potentials (mV) of functionalized AuNPs AuNP@PEG5K, AuNP@PEG5K@R2abF and AuNP@PEG5K@R2abF@ELPA4C. Data points are shown explicitly, and error bars represent the standard error of the mean for at least three independently prepared samples. Statistical significance between grouped samples was determined using one-way ANOVA and Tukey’s multiple comparisons test (****, p < 0.0001).

We needed a rapid assay to assess whether R2ab was accessible and functional when incorporated into nano-assemblies, so we leveraged R2ab’s ability to bind bare polystyrene surfaces, which was first quantified by Somarathne *et al.*.^40^ We hypothesized that R2ab-functionalized TRNs would also bind to bare polystyene,^44^ allowing us to rapidly assess formulations without needing to grow biofilms. A successful formulation would result in nanoparticles that stained bare polysytyene surfaces a light pink color. Here, we added 100 µL of AuNP@PEG5K@R2abF nano-assemblies to polystyrene plates, with each assembly having a different AuNP:PEG5K:R2abF ratio. The ratios ranged from 1:0:250 to 1:225:250, and, as expected, R2ab-functionalized AuNPs uniformly stained the plates a light pink color (Supplementary Material, **Figure S14**). However, nano-assemblies containing only PEG did not stain polystyrene surfaces (Supplementary Material, **Figure S14**, control well). Staining could be evaluated colorimetrically by measuring the extinction at 530 nm in a plate reader (**Figure 1C**). All the R2abF-containing variants had much greater extinction values than AuNP@PEG5K, demonstrating the R2ab domain is able to target AuNPs to bare polystyrene surfaces (and presumably biofilms) but that PEGylation alone cannot (**Figure 1C**, green bar).

The current preferred strategy for integrating and improving the functions of synthesized nanomaterials is multiplex assembly, where multiple functionalizations are employed to enhance particle performance capabilities. Previous work shows that self-assembly or co-assembly of NPs leads to functional improvements. For example, Amarasekara *et al.*, Sun *et al.*, and Lin *et al.* have functionalized the AuNP surface with biological or synthetic polymer coatings to enhance the structural ability as well as to improve the photothermal transduction effect of AuNPs in the NIR region.^25,41,43^ Our previous study conjugated elastin-like polypeptide (ELP) and a second, inert protein to AuNPs, providing a highly tunable and reversible aggregation transition, leading to photothermal responsiveness.^25^ Including the R2abF domain and PEG5K leads to a similar response with a high photothermal conversion efficiency (η). At temperatures above T_t_, the diameter of AuNP@PEG5K@R2abF@ELPA4C increased to several hundred nanometers, as expected for ELP-functionalized particles (**Figure 1D**). Below the T_t_, the particles readily dispersed. Moreover, T_t_ was tunable depending on the AuNP:ELP ratio, as observed previously (**Table 1**). Investigations of aggregation reversibility revealed an unexpected trend in the aggregated D_H_, where the high-temperature D_H_ decreased with each cycle (**Figure 1D**, cycle 1 vs. cycle 5). This could be due to structural changes that occur in nano-assemblies throughout cooling and heating cycles. To probe the source of this behavior, we compared UV-vis extinction spectra between freshly prepared AuNP@PEG@R2abF@ELPA4C and samples that had been thermally cycled five times (Supplementary Material, **Figure S10**). No difference is seen in the plasmonic peak, suggesting the trend in D_H_ (**Figure 1D**) may originate from protein structural changes as opposed to subtle differences in nanoparticle agglomeration. The lower accuracy of DLS when monitoring large aggregates may also contribute to this trend.^54^

**Table 1.** Effect on ELP: AuNP ratio on T_t_ of Targeted TRNs. The AuNP:PEG:R2abF ratio was held constant at 1:125:250.

| AuNP:ELP Ratio | Transition Temperature<br>($T_t$, °C) |
| --- | --- |
| 1:0 | - |
| 1:125 | 42.0 ± 0.1 |
| 1:250 | 39.3 ± 0.6 |
| 1:375 | 39.0 ± 0.1 |
| 1:500 | 38.3 ± 0.6 |
| 1:675 | 38.3 ± 0.6 |

Next, we wanted to determine if the addition of ELP reduced the ability of R2abF to bind surfaces. We used the same colorimetric assay described above with targeted TRN constructs containing AuNP@PEG5K@R2abF@ELPA4C. To start, we selected the construct with the most favorable properties described above (**Table 1**) and chose the R2abF:ELPA4C ratio of 1:1, which had T_t_ that was slightly elevated relative to physiological temperature (39 °C). Again, the colorimetric assay saves time and demonstrates that ELP and R2abF are compatible when functionalized on the same nanoparticle, and the targeted TRN is able to stain polystyrene surfaces more strongly than a control lacking R2abF (**Figure 1E**). This result strongly suggests that the R2ab functionality is retained on these engineered nanospheres containing PEG, ELP, and R2abF. The targeted TRNs were further characterized using DLS, TEM, UV-vis, and zeta potential to examine the sequential addition of each component to the AuNP surface. The addition of PEG5K, R2abF, and ELPA4C to AuNPs resulted in an increase in D_H_ (**Figure 1F**), but no evidence of aggregation below T_t_ was observed via TEM (Supplementary Material, **Figure S7**). The localized plasmon resonance peak of AuNPs at 520 nm redshifts slightly by 2 nm (**Figure 1G**), likely arising from the binding of thiol groups and proteins in our constructs.^25,30^ Finally, the zeta potential increased after adding each component (**Figure 1H**), corresponding to PEG5K binding, then R2abF, and finally ELPA4C.^55^ While the lower magnitude of the zeta potential indicates reduced colloidal stability, we experienced no issues with unexpected aggregation and could readily concentrate the particles to 300 nM (3,600 ppm Au).

### Colloidal stability and cytotoxicity of targeted TRNs

Nanoparticles frequently interact with a variety of biological fluids containing a wide range of proteins and ions that vary significantly from one individual to the other. The layer of proteins that forms is called a biological corona, and it can interfere with nanoparticle targeting.^21,56,57^ We sought to determine whether the targeted TRNs interact with blood proteins to form a stable corona that could block biofilm targeting. D_H_ was measured for citrate AuNPs, AuNP@PEG5K, AuNP@GB3NF, AuNP@ELPA4C, AuNP@PEG5K@R2abF, and AuNP@PEG5K@R2abF@ELPA4C with and without 30% fetal bovine serum (FBS, **Figure 2A**). A significant protein corona forms on non-functionalized AuNPs as well as AuNPs functionalized with GB3NF and R2abF alone. However, the addition of ELPA4C appears to passivate the particles, and no significant increase in D_H_ was observed for targeted TRNs containing all three components. Similar behavior was observed for AuNP@PEG5K and AuNP@ELPA4C nano-assemblies. Thus, while R2abF and GB3NF alone appear to drive corona formation, the inclusion of PEG5K and ELP counteracts this behavior, giving targeted TRNs good surface passivation and reducing corona formation.^58,59^

**Figure 2.**
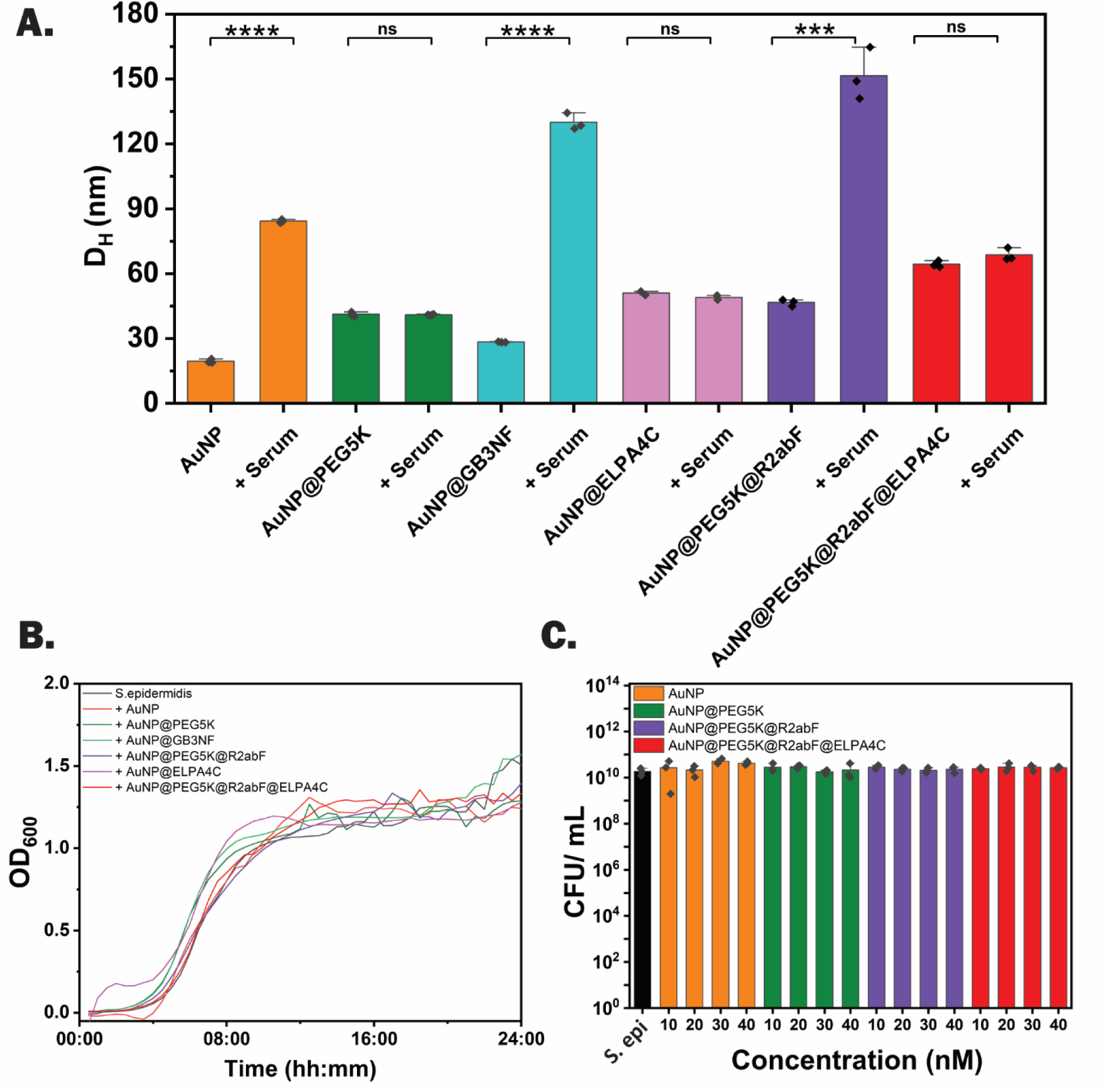
Colloidal stability and cytotoxicity analysis of nano-assemblies. (A.) Observed hydrodynamic diameter (D_H_) of nano-assemblies (AuNP, AuNP@PEG5K, AuNP@GB3NF, AuNP@ELPA4C, AuNP@PEG5K@R2abF, AuNP@PEG5K@R2abF@ELPA4C) with and without 30% (v/v) fetal bovine serum. (B.) Growth curves of *S. epidermidis* bacteria in the presence of 20 nM (120 ppm of Au) AuNPs. (C.) Effect of different concentrations of nano-assemblies on *S. epidermidis* cell growth measured by counting the colony forming units (CFU mL^-1^). Significance in panel A was assessed using one-way ANOVA with Tukey’s multiple comparisons test (ns, not significant; ***, p < 0.001; ****, p < 0.0001). Error bars represent the SEM of three experiments, and all data points are shown.

We next tested the targeted TRNs to see if they affected the viability of *S. epidermidis* bacterial cells. Previously, we found that TRNs lacking R2ab did not affect bacterial growth rates, nor did they alter the viability of human HEK-293 cells grown in cell culture.^25^ As expected, the targeted TRNs containing R2ab did not alter our previous results. A final AuNP concentration of 20 nM (240 ppm total Au) was added to a fresh culture of S. *epidermidis,* and the cell growth curve was monitored using a plate reader (**Figure 2B**). All the nano-assemblies used in the study showed no change in cell growth compared to the control, confirming that all the nano-assemblies (including targeted TRNs) are biocompatible at an Au concentration of 240 ppm after incubation for 24 h. This is consistent with prior studies of functionalized AuNPs; when compounds such as cetyl-trimethylammonium bromide (CTAB) are avoided, functionalized AuNPs are generally biocompatible.^60–62^ Even at 40 nM (480 ppm Au), no observable effect on bacterial growth was observed for any of the constructs tested (**Figure 2C**). This biocompatibility extends to mammalian cells as well. Cell viability studies using HEK-293 cells showed all nanoparticle formulations maintained >90% cell viability even at 40 nM concentration (Supplementary Material, **Figure S18**), further supporting the safety profile of these constructs. Thus, in the absence of NIR irradiation, the nano-assemblies used in this study resist corona formation and do not adversely affect cell function.

### Assessing selective biofilm adherence and penetration of targeted TRNs

Next, we sought to assess the effectiveness of bacterial targeting using the engineered R2ab fusion domain. R2ab targets the cell wall components of *S. epidermidis*,^28^ and we hypothesized that the R2ab function would localize the targeted TRNs to *S. epidermidis* biofilms, enhancing their effect. To start, we tested the interaction with biofilms grown statically in a 96-well plate. Biofilms were treated with 20 nM (240 Au ppm) of AuNP@PEG5K@R2abF@ELPA4 with or AuNP@PEG5K as a control (see Materials and Methods). Biofilms were washed three times to remove excess TRNs, and the remaining nanoparticles were measured using extinction at 530 nm (Supplementary Material, **Figure S15**). The retention of all targeted TRNs (regardless of ELP content) was significantly higher than the AuNPs coated with PEG alone. To assess binding more quantitatively, we digested biofilms in aqua regia and measured total content using inductively coupled plasma-mass spectrometry (ICP-MS, **Figure 3A**). Of the six systems tested, only targeted TRNs containing ELP, PEG5K, and R2abF showed the desired specificity, where adhesion to biofilms was higher than to serum-coated surfaces. Interestingly, AuNPs lacking ELP (but containing R2ab) had the opposite effect, where the particles bound more favorably to serum-coated polystyrene surfaces. This behavior likely results from ELP’s anti-fouling properties, which prevents unwanted nonspecific binding to serum (**Figure 2A**). For all the other particles tested, binding to serum-coated surfaces was higher than binding to biofilms, and the lack of any functionalization on citrate AuNPs led to the highest binding of all (**Figure 3A**, orange bars). Thus, targeted TRNs containing PEG5K, ELP, and R2abF exhibit specific binding toward *S. epidermidis* biofilms as designed.

**Figure 3.**
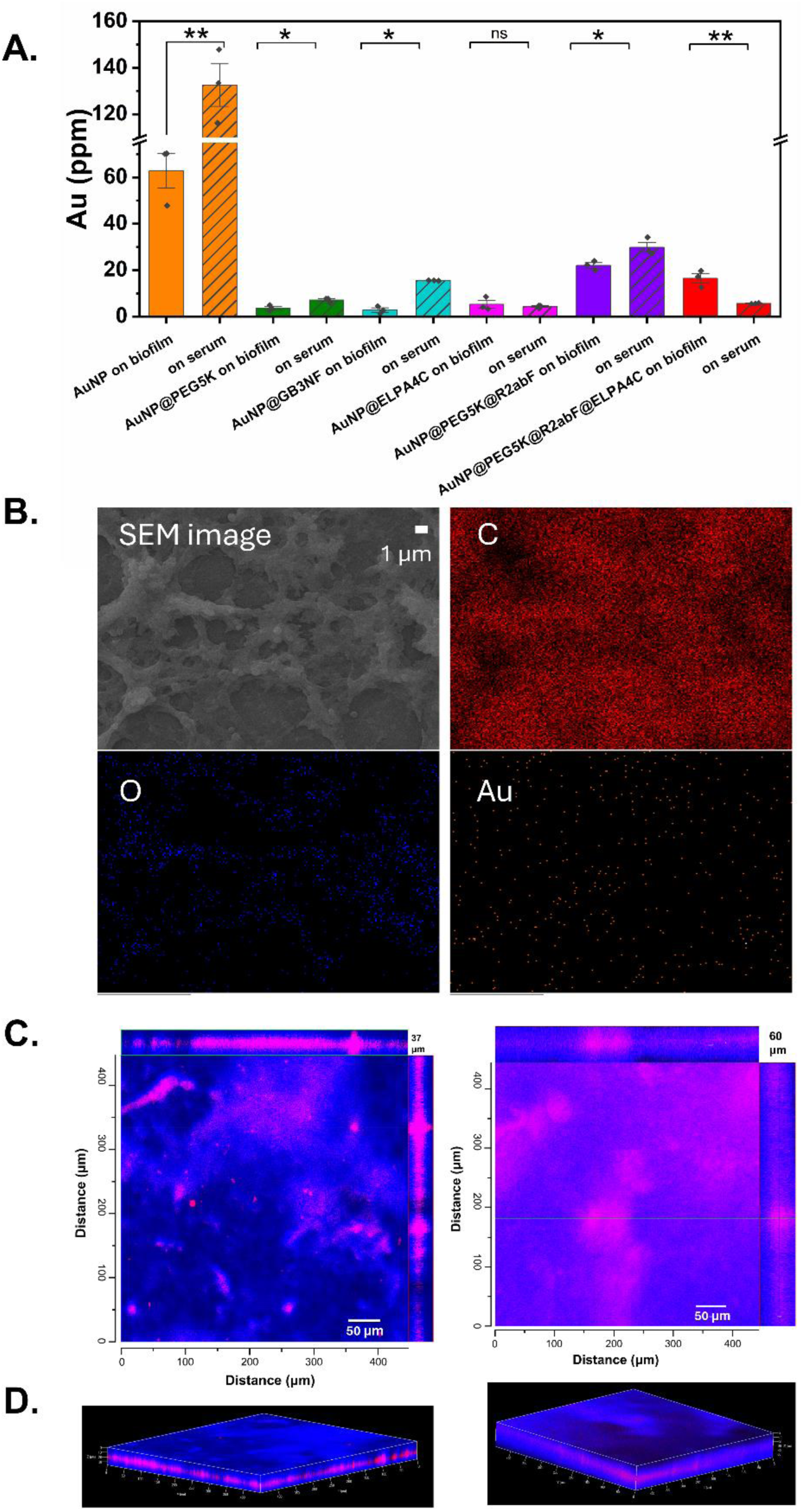
Determining the adherence and penetration of targeted TRNs on different biological surfaces. (A.) Affinity profiles of nano-assemblies on biofilm and serum-coated polystyrene surfaces measured using ICP-MS. (B.) SEM images of *S. epidermidis* biofilm formed on polystyrene surface treated with AuNP@PEG5K@R2abF@ELPA4C and analyzed with EDS elemental analysis. EDS indicates the presence of carbon (C), oxygen (O), and gold (Au) on the surface of the treated biofilms. (C.) Orthogonal views of confocal microscopy images showing the penetration of Texas Red-labeled AuNP@PEG5K@R2abF@ELPA4C nanoparticles in an *S. epidermidis* biofilm. (D.) Three-dimensional reconstructions of confocal microscopy Z-stack data demonstrating the distribution of targeted nanoparticles throughout the biofilm structure. Significance in panel A was assessed using one-way ANOVA with Tukey’s multiple comparisons test (ns, not significant; *, p < 0.05; **, p < 0.01). Error bars represent the SEM of three experiments and all data points are shown.

To visualize specific binding, we used scanning electron microscopy (SEM) coupled with energy dispersive X-ray spectroscopy (EDS) to monitor the presence of targeted TRNs (but not AuNPs@PEG5K) on biofilm surfaces. The characteristic peak at 2 keV confirmed the presence of gold on the biofilm surface when treated with targeted TRNs; the corresponding peak had a much lower relative intensity when biofilms were treated with PEGylated AuNPs alone (Supplementary Material **Figures S8** and **S9**). Visually, the Au signals from EDS appear localized to regions where biofilm is present and not the polystyrene background (**Figure 3B** and Supplementary Material, **Figure S16**). Thus, optical staining, quantitative measurement by ICP-MS, and electron microscopy all support that targeted TRNs containing the *S. epidermidis* R2ab domain are able to bind *S. epidermidis* biofilms selectively, whereas nanoparticles lacking R2ab do not.

We also used confocal fluorescence microscopy to visualize the penetration of targeted TRNs into biofilms. Quantitative analysis of confocal Z-stack data revealed that *S. epidermidis* biofilms (shown in blue) grown on cell culture slides achieved an average thickness of 46.7 ± 7.9 μm. The Texas Red-labeled AuNP@PEG5K@R2abF@ELPA4C nanoparticles (shown in magenta) possessed substantial penetration capability, reaching approximately 60% of the total biofilm thickness, corresponding to an average penetration depth of ∼28 μm as seen in three-dimensional reconstructions of the Z-stack data (**Figure 3C, D**). This deep penetration suggests that the R2ab targeting domain enables the nanoparticles to effectively navigate through the complex extracellular polymeric matrix and reach bacterial cells embedded within the biofilm structure. It also suggests that binding to biofilms enhances the local concentration of targeted TRNs, which will reduce the T_t_, enhancing the photothermal conversion efficiency (η).^25^ The penetration profile showed relatively uniform distribution across the biofilm thickness, with targeted nanoparticles observed to accumulate in regions of high bacterial density throughout the biofilm depth.

Similar penetration characteristics were observed with *S. aureus* biofilms (Supplementary Material, **Figure S17**), which exhibited an average thickness of 60.0 ± 5.0 μm but showed a similar nanoparticle penetration of approximately 60% of the total biofilm thickness (corresponding to ∼36 μm penetration depth). The consistency of penetration performance across both staphylococcal species, despite differences in biofilm architecture and thickness, suggests that the R2ab targeting mechanism is broadly applicable for biofilm-associated infections, including methicillin-resistant *S. aureus* (MRSA). This is because S. aureus cell walls also contain substantial LTA and WTA, which are the targets of R2ab.^63^

### In vitro antibiofilm activity of targeted TRNs under NIR irradiation

Next, we sought to assess the effectiveness of biofilm-targeted TRNs in eradicating biofilm bacteria upon exposure to NIR irradiation. Solutions of targeted TRNs were added to *S. epidermidis* biofilm-coated polystyrene surfaces and then exposed to NIR light to assess the bactericidal impact. Constructs lacking ELP and R2abF were used as controls. After being exposed to NIR light for the first five minutes, the *S. epidermidis* biofilm containing AuNPs all show an increase in the temperature (**Figure 4A-C**). However, at or above 20 nM particle concentration (240 ppm of Au), the targeted TRNs outperformed the temperature increase of the controls. At 30 nM (360 ppm of Au), a maximum temperature of around 60 °C is obtained, which is sufficient to kill bacteria embedded in biofilms.^25^ The photothermal conversion efficiency (η) of AuNP@PEG5K@R2abF@ELPA4C was calculated to be around 40 ± 2% (Supplementary Material, **Figure S11**), which was higher than that of AuNP@PEG5K (31 ± 5%) as reported previously.^25^ After treatment with targeted TRNs and exposure to NIR light for 5 min, the number of surviving bacteria dramatically decreased (**Figure 4D** and **4E**), demonstrating an enhanced photothermal bactericidal effect that was absent for TRNs lacking ELP. Targeted TRNs aggregate reversibly at elevated temperatures, which leads to this enhanced photothermal effect (Supplementary Material, **Figure S12**).

**Figure 4.**
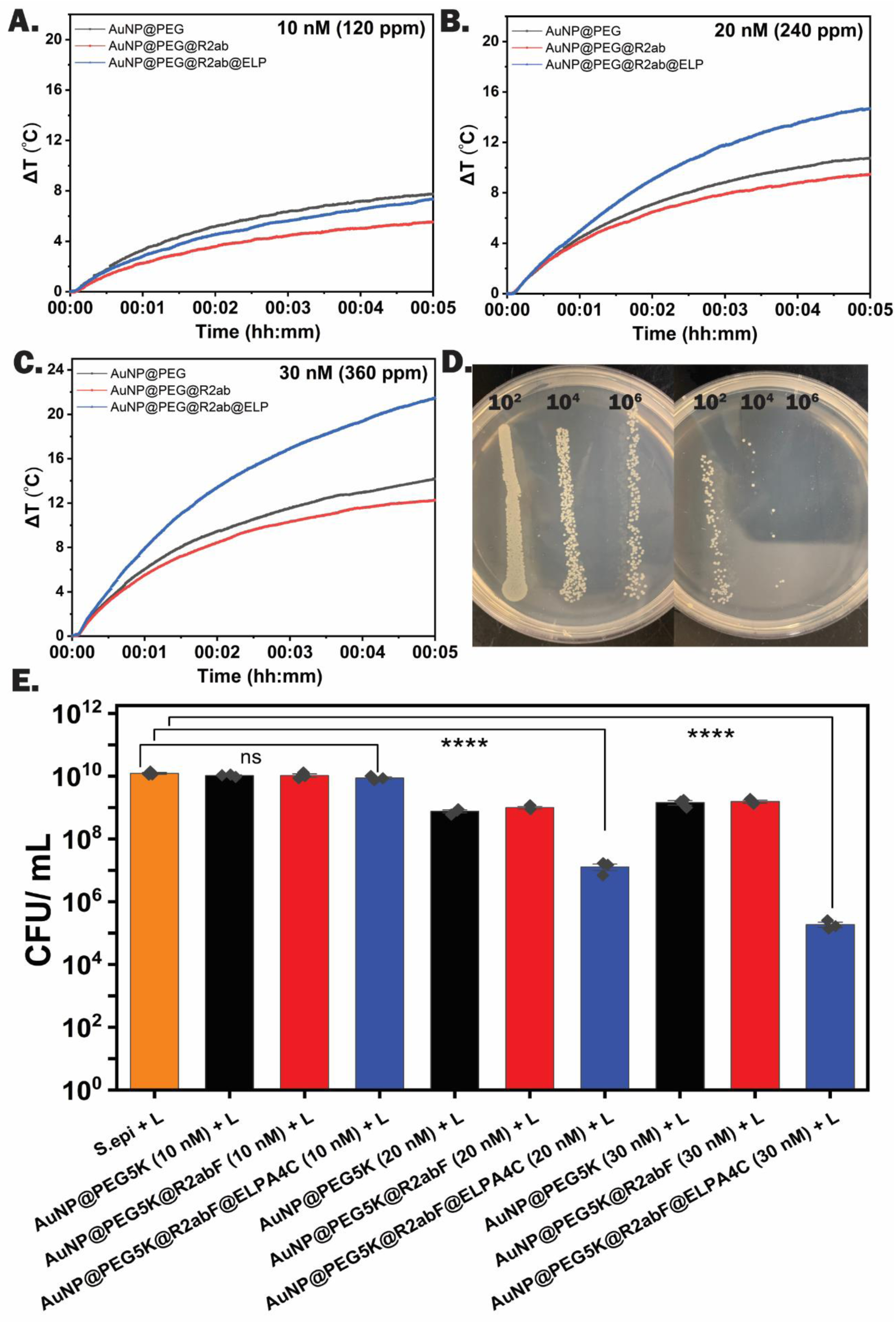
Determining the anti-bacterial ability of AuNP@PEG5K@R2abF@ELPA4C. Temperature evolution curves of (A.) 10 nM (B.) 20 nM (C.) 30 nM AuNP@PEG5K, AuNP@PEG5K@R2abF and AuNP@PEG5K@R2abF@ELPA4C added to *S. epidermidis* biofilms under NIR Irradiation above the transition temperature (T_t_). (D.) Serial dilution assay on agar plates demonstrating how CFU mL^-1^ is determined. (E.) CFU mL^-1^ of *S. epidermidis* present in biofilms after treatment with targeted TRNs at 10-30 nM particle concentrations. AuNP@PEG5K and AuNP@PEG5K@R2abF were used as controls. Significance was assessed using one-way ANOVA with Tukey’s multiple comparisons test (n.s., not significant; ****, p < 0.0001). Error bars represent the SEM of three experiments, and all data points are shown.

### Assing the anti-biofilm effectiveness of targeted TRNs under continuous flow conditions

The experiments above demonstrate that targeted TRNs possess the same photothermal killing as our original TRNs, which lack a biofilm targeting domain;^25^ however, the additional PEG and R2abF confer a useful biofilm-targeting function. It remains to test whether the targeting domain is sufficient to localize TRNs to biofilms in the presence of continuous flow, which is more similar to what is found in in vivo systems. An inexpensive dynamic biofilm reactor was developed to study nanoparticle adhesion properties under continuous flow through polystyrene tubing derived from a sterile serological pipet. We used a 3D printed base (**Figure 5A** and **5B**) to hold 3 tubes in series. A 12 V peristaltic pump was used to drive a flow of 5 mL min^-1^ (linear rate of 4.6 cm s^-1^) of BHI medium and targeted TRNs through the tubes. For the 1.5 mm inner diameter tubes used here, this rate is comparable to what is experienced in the arteries of the circulatory system.^27,64,65^ Our system is similar to the Modified Robbins Device (MRD) discussed by Coenye and Nelis.^66^ Polytetrafluoroethylene (PTFE) tubing was used to circulate the solution, as biofilm growth was significantly reduced on PTFE surfaces. New PTFE tubing and polystyrene tubes were used after each experiment to maintain consistency between experiments. The plans for the 3D-printed components of the biofilm flow reactor have been included in the online Zenodo repository (see Materials and Methods).

**Figure 5.**
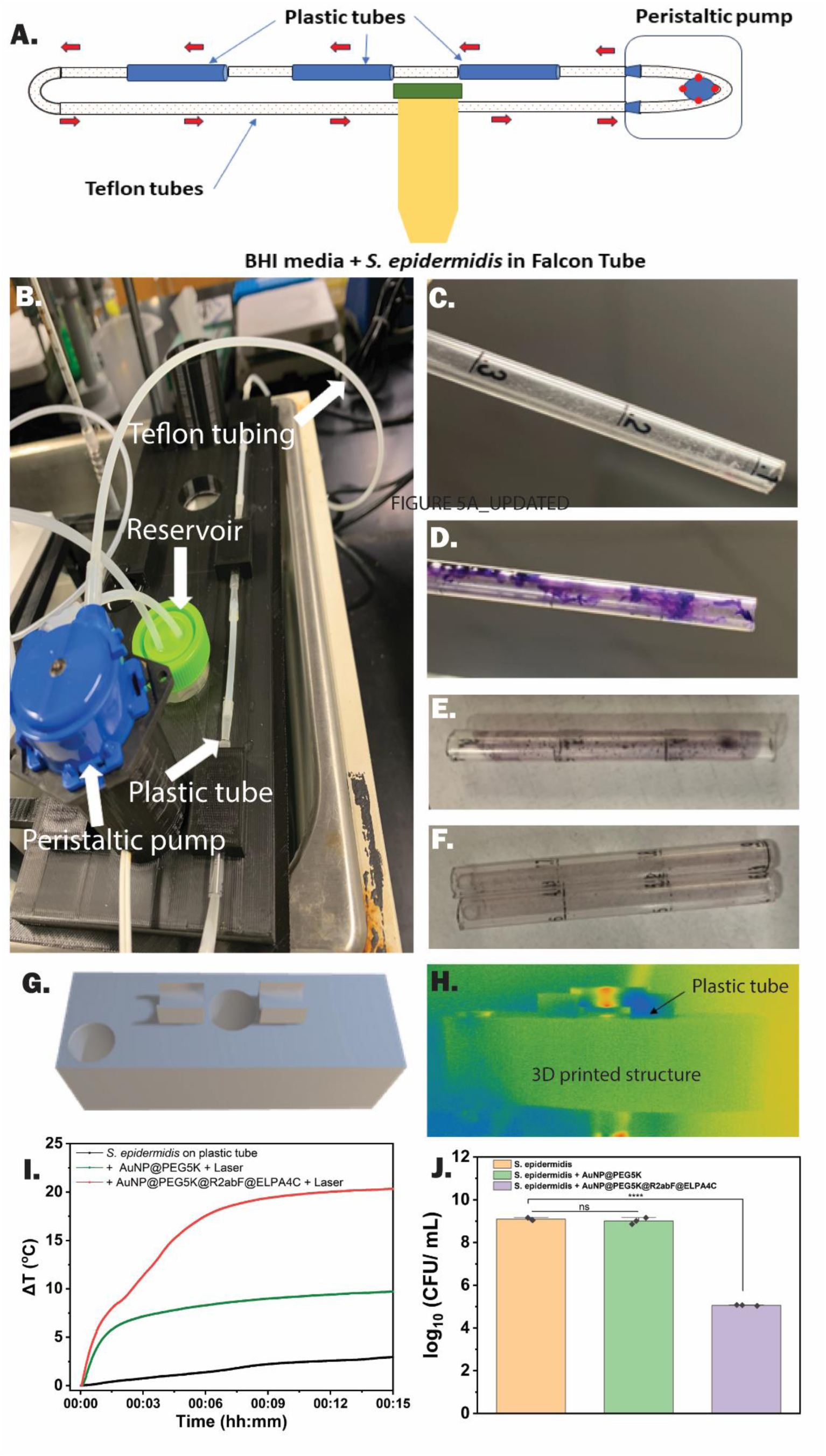
Determining the anti-bacterial ability of AuNP@PEG5K@R2abF@ELPA4C in a dynamic flow system. (A.) Schematic diagram (B.) Actual image of the dynamic flow system. (C.) Visible biofilm formation on plastic tubes (3 cm) after flowing BHI media containing *S. epidermidis* cells for 48 h. (D.) Crystal violet staining to determine the presence of *S. epidermidis* biofilm on the inner walls of plastic tube (E.) Presence of AuNP@PEG5K@R2abF after flowing 20 nM AuNP@PEG5K@R2abF through the biofilm containing tubes for 12 h. (F.) Presence of AuNP@PEG5K@R2abF@ELPA4C after flowing 20 nM AuNP@PEG5K@R2abF through the biofilm containing tubes for 12 h. (G.) Image of 3D printed structure used to measure the temperature evaluation curves of plastic tubes under the NIR irradiation. (H.) Thermal camera image and (I.) Temperature evaluation curves of plastic tubes after irradiation with 1.8 W cm^-2^ NIR laser for 15 min. (J.) Colony formation after the treatment with NIR laser. Significance was assessed using one-way ANOVA with Tukey’s multiple comparisons test (n.s., not significant; ****, p < 0.0001). Error bars represent the SEM of three experiments, and all data points are shown.

By using this platform to conduct 48 h dynamic biofilm tests with *S. epidermidis*, we were able to detect biofilm formation on the inner walls of the plastic tubes (**Figure 5C**). Crystal violet staining (**Figure 5D**) and SEM imaging (Supplementary Material, **Figure S13**) were used to confirm the presence of biofilm. The excess unbound bacteria on the inner walls of the plastic tubes were removed by washing with PBS buffer. AuNPs, AuNP@PEG5K, and AuNP@PEG5K@R2abF@ELPA4C (30 nM or 360 Au ppm) were circulated through the system to assess the biofilm adhesion on the plastic tubes separately. Despite the presence of flow, our results were consistent with the experiments performed under static conditions (**Figure 3A**). No biofilm binding was detected for AuNP@PEG5K particles, but the addition of the R2ab fusion domain (AuNP@PEG5K@R2abF) effectively stained the biofilm red (**Figure 5E**). The complete targeted TRN particle (AuNP@PEG5K@R2abF@ELPA4C) also stained biofilms (**Figure 5F**), demonstrating its ability to adhere to the biofilm even under dynamic conditions.

Next, we determined the photothermal properties of the biofilm-bound nano-assemblies using an 808 nm NIR laser. A 3D-printed structural mount was used to hold biofilm-coated tubes during laser irradiation (**Figure 5G**). After treatment with nanoparticles, each tube was cut into three 1 cm sections. These sections, containing biofilm only, biofilm and AuNP@PEG5K, or biofilm and targeted TRNs (AuNP@PEG5K@R2abF@ELPA4C), were irradiated with 1.8 W cm^-2^ 808 nm NIR laser for 15 min to observe their temperature response (**Figure 5H**). A longer laser treatment time was needed to reach a steady temperature, presumably due to the decreased concentration of TRNs. However, a significant temperature increase was obtained for targeted TRNs (**Figure 5I**), where the final temperature increase was nearly 13 °C more than AuNP@PEG5K (red vs. green curve). To assess the bactericidal effect, each 1 cm section was soaked in 1 mL of PBS media with agitation for 30 min, allowing the live cells to detach from the biofilm. The CFU mL^-1^ of this solution was measured using the serial dilution assay. Significantly fewer cells grew from tubes treated with targeted TRNs and NIR irradiation than the untreated control or tubes treated with AuNP@PEG5K and NIR irradiation (**Figure 5J**); an almost 10^4^-fold reduction in CFU mL^-1^ was observed. This was consistent with bacteria grown in static biofilms (**Figure 4E**). Thus, the targeted TRNs developed here effectively target photothermally active nanoparticles to *S. epidermis* biofilms, both under static and flow conditions. This work demonstrates that bacterial proteins such as R2ab, which bind cell wall components, can be used to functionalize nanomaterials, effectively targeting those nanomaterials to infection sites.

## Conclusions

Recently, it was shown that nanoparticles functionalized with Hoechst dye could be used to target nanoparticles to biofilms, since this dye can form a specific adduct with extracellular DNA in the biofilm.^67^ Here, we employ a similar strategy with a domain derived from the bacteria’s own extracellular autolysin. Using the *S. epidermidis* R2ab domain, we can target nanoparticles to *S epidermidis* biofilms. The mechanism is driven by R2ab’s natural ability to bind LTA and WTA cell wall components, which are not expected to be present in mammalian tissues. We synthesized temperature-responsive, bacterial biofilm targeting spherical AuNPs using cystine-based conjugation and investigated their photothermal properties, including their performance in a simple anti-biofilm assay. These targeted TRNs (AuNP@PEG5K@R2abF@ELPA4C) agglomerate in response to NIR irradiation, forming particle clusters with high photothermal conversion efficiency. Effective and selective bactericidal properties were confirmed for both statically grown biofilms and biofilms grown and treated under continuous flow. The temperature responsiveness and selective adhesion of the synthesized particles can potentially be employed for developing intelligent photothermal transducers for non-surgical thermal ablation of *S. epidermidis* biofilms. The targeted TRNs optimized here provide a starting point for in vivo studies, where biofilms could be treated non surgically using photothermal therapy.

## ^i^ Abbreviations

AuNP: gold nanoparticle;
DLS: dynamic light scattering;
ELP: elastin-like polypeptide;
EPS: extracellular polymeric substances;
ICP-MS: inductively coupled plasma-mass spectrometry;
LTA: lipoteichoic acid;
NIR: near infrared;
PDT: photodynamic treatment;
PGN: peptidoglycan;
PTA: photothermal agent;
PTT: photothermal therapy;
TEM: transmission electron microscopy;
TRN: temperature-responsive nanosphere;
T_t_: transition temperature;
WTA: wall teichoic acid.

## Authorship Contribution Statement

**Dhanush Amarasekara:** Conceptualization, Data curation, Formal analysis, Investigation, Methodology, Validation, Visualization, Writing – original draft, Writing - review & editing. **Radha Somarathne:** Investigation, Methodology. **Tanveer Shaikh:** Investigation, Methodology, Visualization. **Gabriel J. Alcantara:** Investigation, Methodology. **Madison Hejny:** Investigation, Methodology. **Elizabeth McCaffrey:** Investigation, Methodology. **Nicholas Fitzkee:** Conceptualization, Data curation, Formal analysis, Funding acquisition, Investigation, Methodology, Project administration, Supervision, Validation, Visualization, Writing – original draft, Writing - review & editing.

## Data Availability

All data have been uploaded to Zenodo and are available at https://doi.org/10.5281/zenodo.10887304.

## Acknowledgments

This work was supported by the National Institutes of Health under award number R56AI139479 (NCF) and the National Science Foundation under awards OIA 2414443 (NCF), CBET 2405018 (NCF), OIA 2414442 (TAW), and the American Cancer Society under award RSG-21-114-01-MM (TAW). Support for the MSU NMR Facility was provided by the National Science Foundation under awards CHE/MCB 2304919 and DBI 2215258.

## Supplementary Material

Supplementary data for this article can be found online at …

