## Supplemental Material for "Using a Bacterial Protein to Selectively Target Bacterial Biofilms: Treatment of *S. epidermidis* Biofilms with Targeted Photothermal Gold Nanoparticles"

**Supporting Figures**

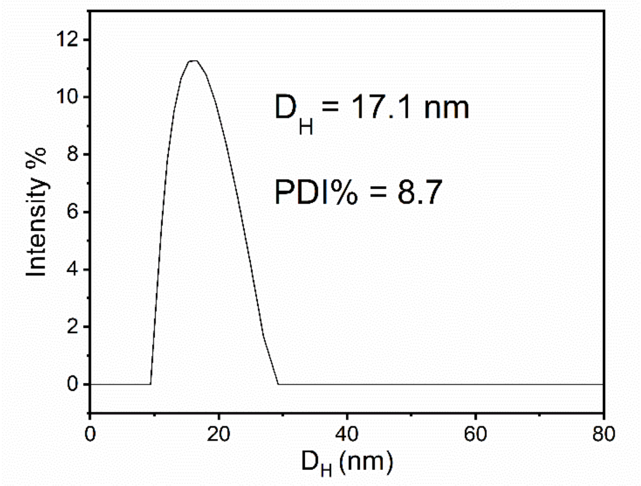

**Figure S1. Size distribution of synthesized AuNPs.**

Particle size distribution (PSD) of 15 nm citrate AuNPs was measured using an Anton Paar Dynamic Light Scattering (DLS) system at room temperature. The average hydrodynamic diameter ($D_H$) and polydispersity index (PDI%) are shown.

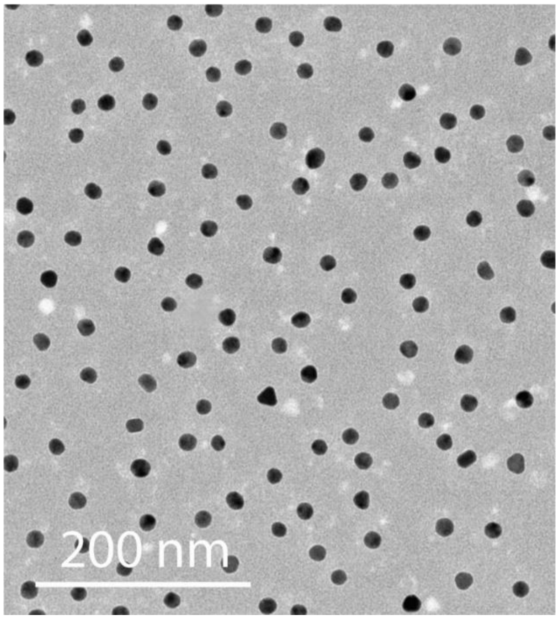

**Figure S2. TEM image of synthesized AuNPs.**

Transmission electron microscopy (TEM) image of 15 nm synthesized citrate AuNPs shows a uniform size distribution (imaged using a JEOL 2100 TEM instrument)

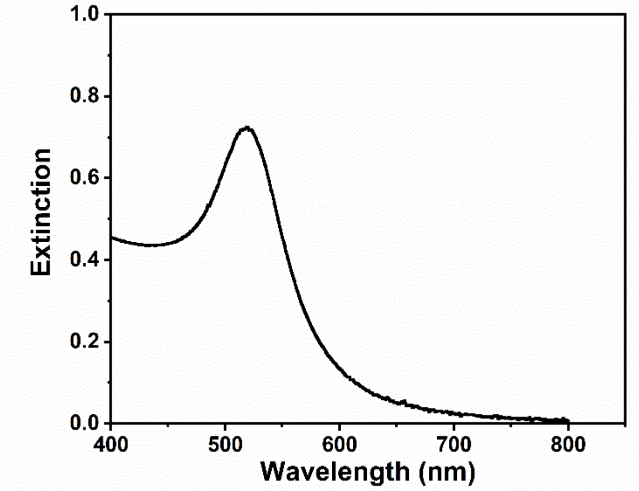

**Figure S3. Extinction spectrum of AuNPs.**

UV-vis extinction of synthesized 15-nm AuNPs as a function of wavelength was measured using an Olis-refurbished, Peltier-controlled Agilent 8453 UV-vis spectrophotometer. The single peak at 520 nm indicates uniform, spherical AuNPs with a diameter of approximately 15 nm.

```
GSSMQYKLVI CGKTLKGETT TKAVDAETAE KAFKQYANDN GVDGVWTYDD
ATKTFTVTEG GGGSGSAGSE AAGSEGSAGS EAAGSEGGGG SENLYFQSGS
HMSSTNNQLT VTNNSGVAQI NAKNSGLYTT VYDTKGKTTN QIQRTLSVTK
AATLGDKKFY LVGDYNTGTN YGWVKQDEVI YNTAKSPVKI NQTYNVKPGV
KLHTVPWGTY NQVAGTVSGK GDQTFKATKQ QQIDKATYLY GTVNGKSGWI
SKYYLTA
```

**Figure S4. Amino acid sequence of R2ab fusion protein (R2abF).**

A schematic representation of R2ab fusion sequence is shown. The cysteine residue is highlighted and printed in red text.

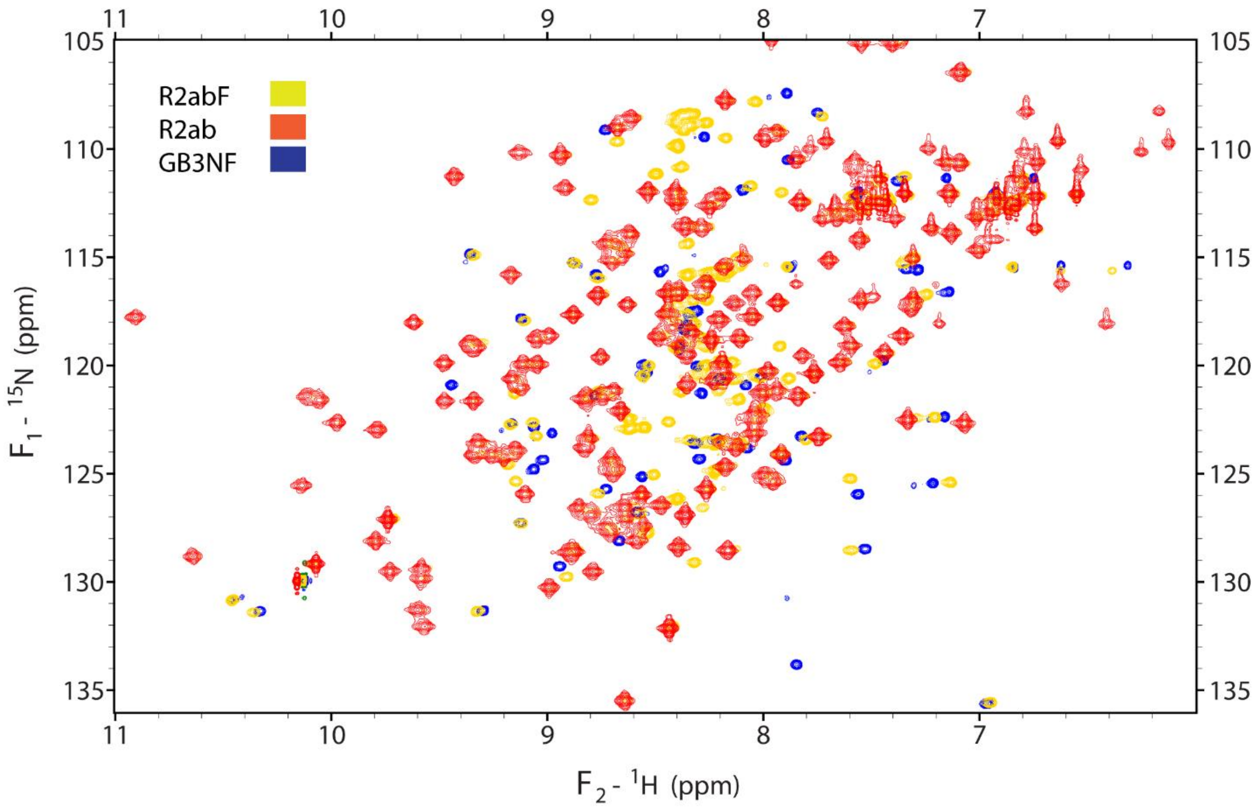

**Figure S5. $^{15}N$–$^{1}H$ TROSY spectrum of R2abF, R2ab, and GB3NF**

2D TROSY spectra were collected for the protein constructs used in this work. An overlay of these spectra of R2abF (gold), R2ab (red) and GB3NF (blue) is shown. No significant shifts are observed for peaks in the R2ab and GB3 domains, indicating that there is no structural perturbation to these domains when connecting them via the flexible linker.

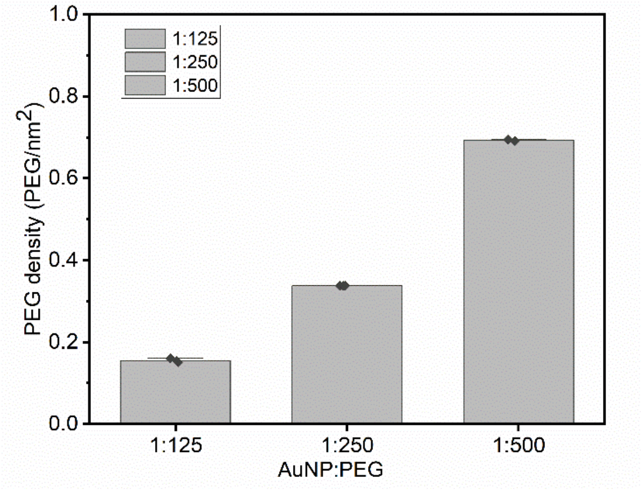

**Figure S6. PEG density on AuNP surface at different AuNP:PEG ratios**

The experimentally measured PEG number density on AuNP-PEG conjugates upon varying the AuNP:PEG ratio, ranging from 0.15 ± 0.01 PEG $nm^{-2}$ at 1:125 to 0.69 ± 0.01 PEG $nm^{-2}$ at 1:500.

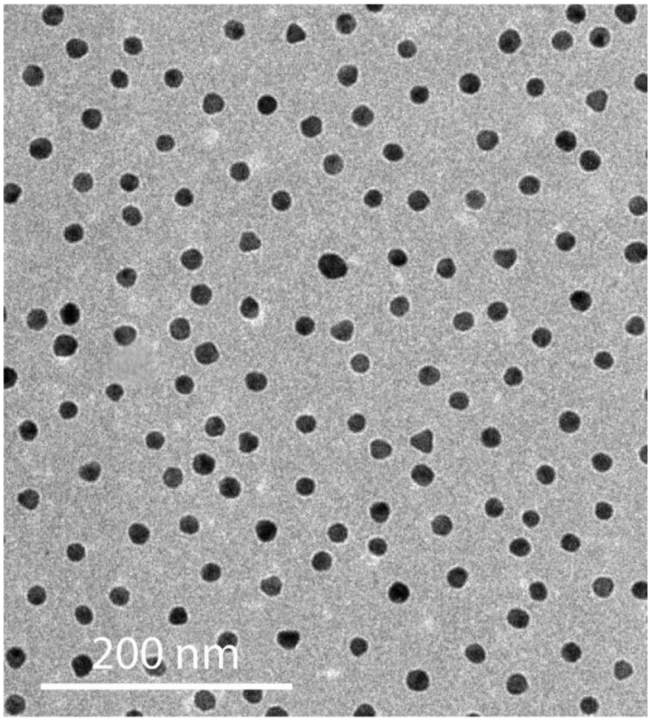

**Figure S7. TEM image of targeted thermally-responsive nanospheres (TRNs).**

Transmission electron micrographs demonstrate that the AuNP@PEG5K@R2abF@ELPA4C TRNs are not aggregated below the transition temperature ($T_t$).

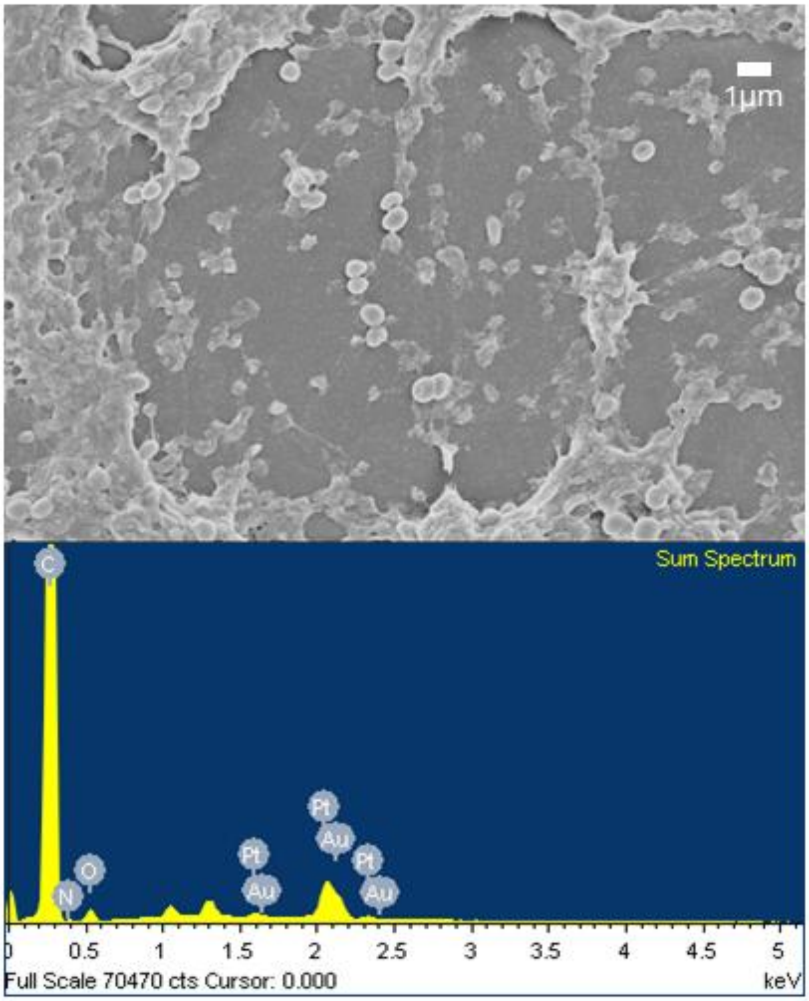

**Figure S8. SEM and EDS of *S. epidermidis* biofilm on polystyrene treated with PEGylated AuNPs**

Scanning electron microscopy (SEM) with energy dispersive X-ray spectroscopy (EDS) was used to visualize an *S. epidermidis* (strain 1301) biofilm after treatment with PEGylated AuNPs and subsequent washing to remove unbound nanoparticles. (A.) The field used for EDS analysis, (B.) Elemental map from EDS, highlighting the absence of significant gold (Au) peaks. Note that Pt appears because it is used in the sputter coating process when preparing SEM samples.

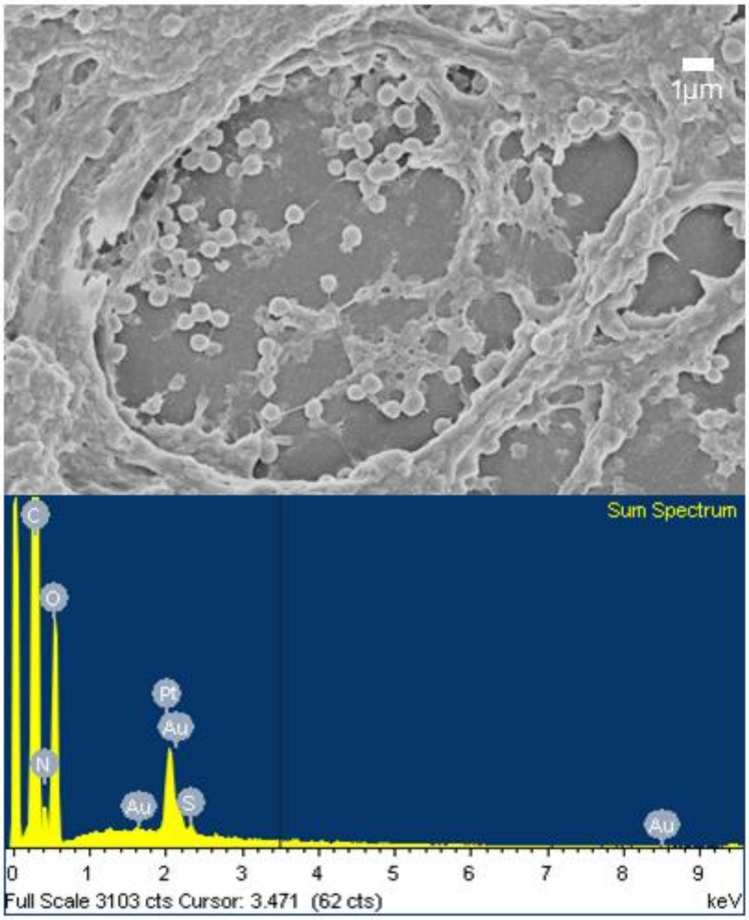

**Figure S9. SEM of *S. epidermidis* biofilms after treatment with targeted TRNs.**

SEM was used to visualize *S. epidermidis* (Strain 1301) as above; here biofilms were treated with AuNP@PEG5K@R2abF@ELPA4C then washed before preparation to identify whether targeted TRNs tightly bind to biofilms. (A.) The field used for EDS analysis, (B.) Elemental map from EDS, highlighting the presence of gold (Au) peaks. Note that Pt appears because it is used in the sputter coating process when preparing SEM samples.

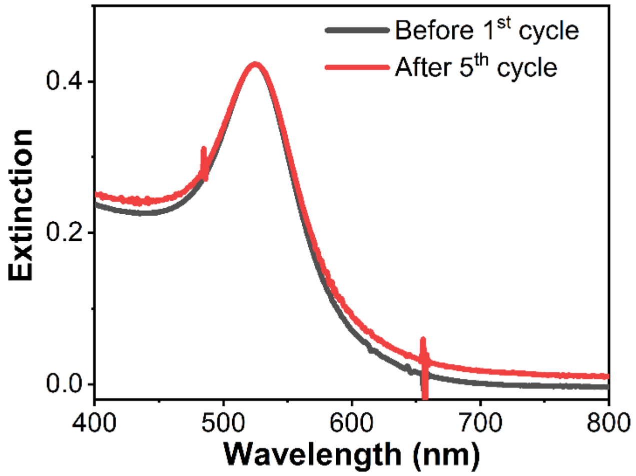

**Figure S10. UV–vis profiles of targeted TRNs before and after heating cycles.**

The UV-vis extinction spectra of AuNP@PEG5K@R2abF@ELPA4C are shown before and after five cycles of heating above $T_t$ and cooling below the $T_t$. The spectra retains its shape after repeated cycles of aggregation.

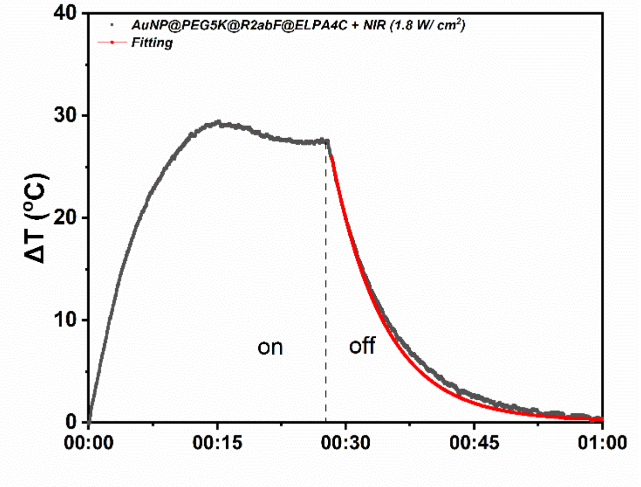

**Figure S11. Photothermal effect of AuNP@PEG5K@R2abF@ELPA4C.**

Sample temperature change with laser irradiation (on) and without (off) for AuNP@PEG5K@R2abF@ELPA4C, when 20 nM (240 ppm Au) aqueous solution was irradiated with a 1.8 W cm$^{-2}$, 808 nm laser at 39 °C. The exponential decay fitting to determine the photothermal conversion efficiency (η) is shown in red.

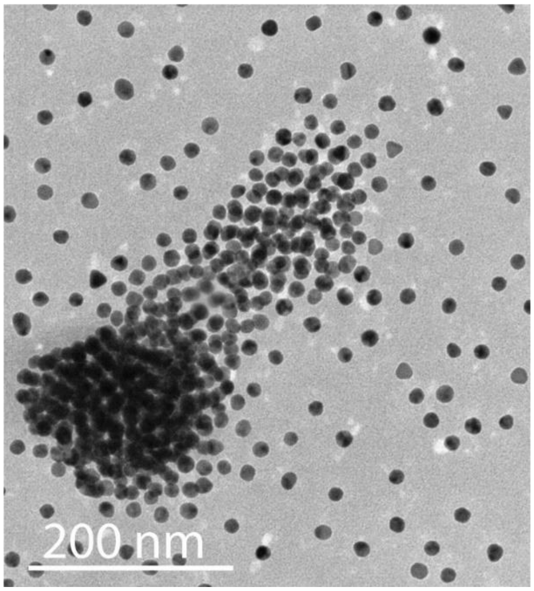

**Figure S12. TEM image of targeted TRNs upon aggregation.**

TEM demonstrates that AuNP@PEG5K@R2abF@ELPA4C are aggregated when prepared above $T_t$. This aggregation leads to enhanced photothermal conversion efficiency of photons in the near-infrared region (NIR).

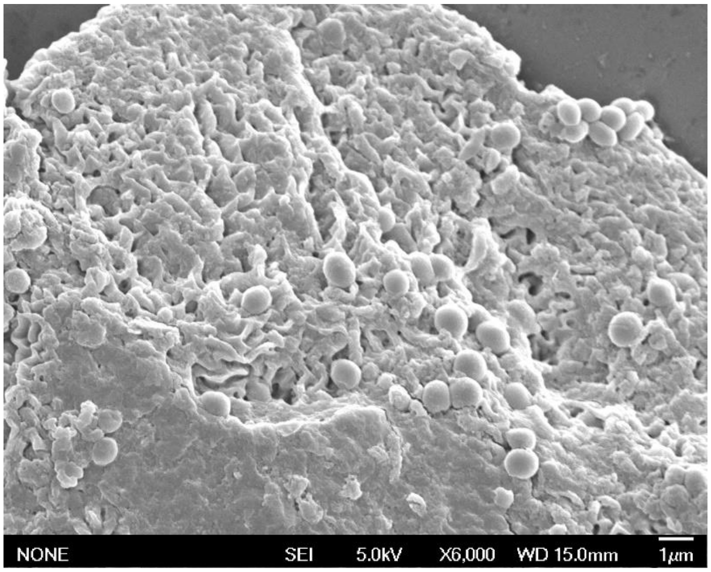

**Figure S13. SEM image of *S. epidermidis* biofilms under dynamic flow conditions.**

After 72 h of continuous dynamic flow through polystyrene tubes, *S. epidermidis* biofilms were collected and visualized by SEM.

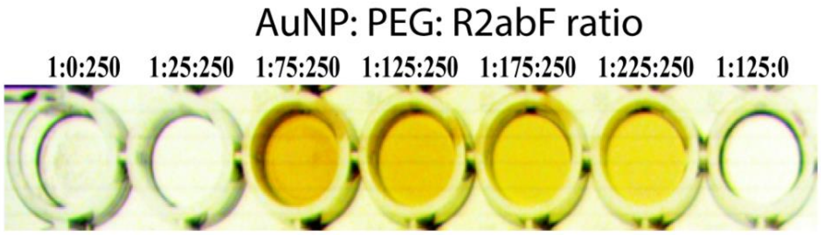

**Figure S14. Affinity of differently functionalized AuNPs with polystyrene surface.**

Image of wells stained with AuNP@PEG5K and AuNP@PEG5K@R2abF, where different ratios of R2abF and PEG5K were used on each AuNP. The brightness and contrast were adjusted to highlight the staining of wells when R2abF is present. The first column shows no staining because AuNP@R2abF rapidly aggregates without prior functionalization with PEG5K; the aggregate readily washes out of the well. The final column shows that AuNPs lacking R2ab do not stain polystyrene at all.

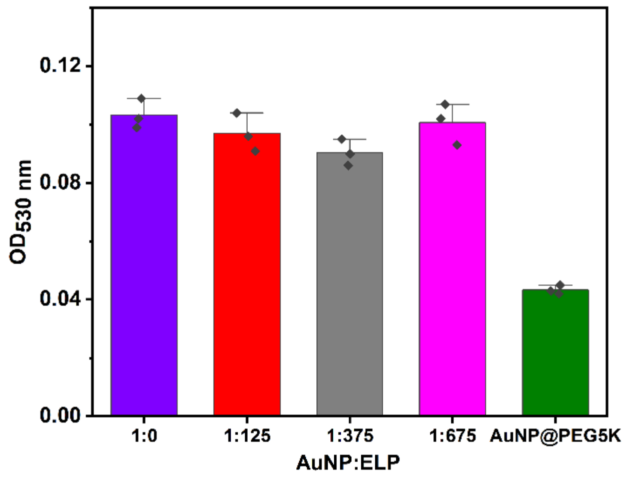

**Figure S15. Selective adhesion of targeted TRNs to *S. epidermidis* biofilms.**

*S. epidermidis* biofilm-binding profiles of AuNP@PEG5K@R2abF@ELPA4C were explored by varying the ratio of elastin-like polypeptide (ELP). After washing, the presence of targeted TRNs was monitored in a plate reader by monitoring the intensity of the pink color (λ = 530 nm). The AuNPs coated with only PEG (green) bind to biofilms least of all.

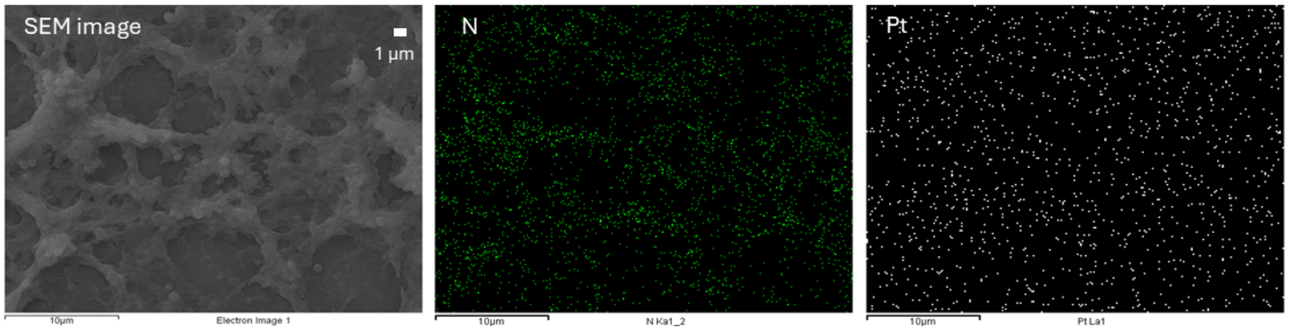

**Figure S16. SEM and EDS of *S. epidermidis* biofilm on polystyrene with targeted TRNs.**

Additional SEM EDS images of *S. epidermidis* biofilm formed on a polystyrene surface are shown after treatment with AuNP@PEG5K@R2abF@ELPA4C. In addition to the images shown in the main text, elemental analysis indicated the presence of Nitrogen (N, from the biological sample), and Platinum (Pt, from sputter coating during sample preparation) on the surface of the biofilms.

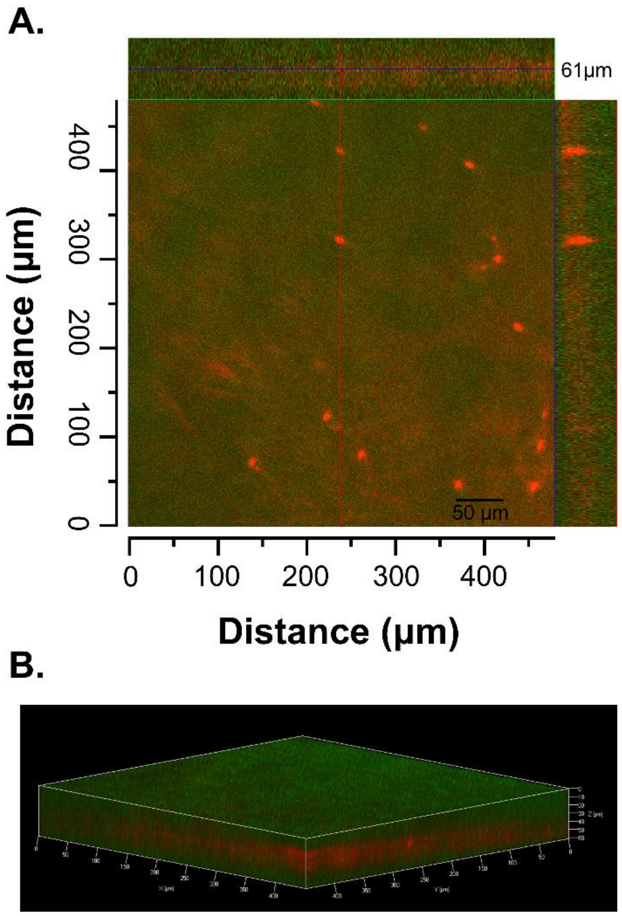

**Figure S17. Confocal microscopy analysis of GFP containing *S. aureus* biofilm treated with Texas Red-labeled AuNP@PEG5K@R2abF@ELPA4C.**

(A) Orthogonal view of Z-stack showing NP penetration (red) through the biofilm (green). Scale bar: 50 μm. (B) 3D reconstruction demonstrating the distribution of nanoparticles throughout the biofilm structure.

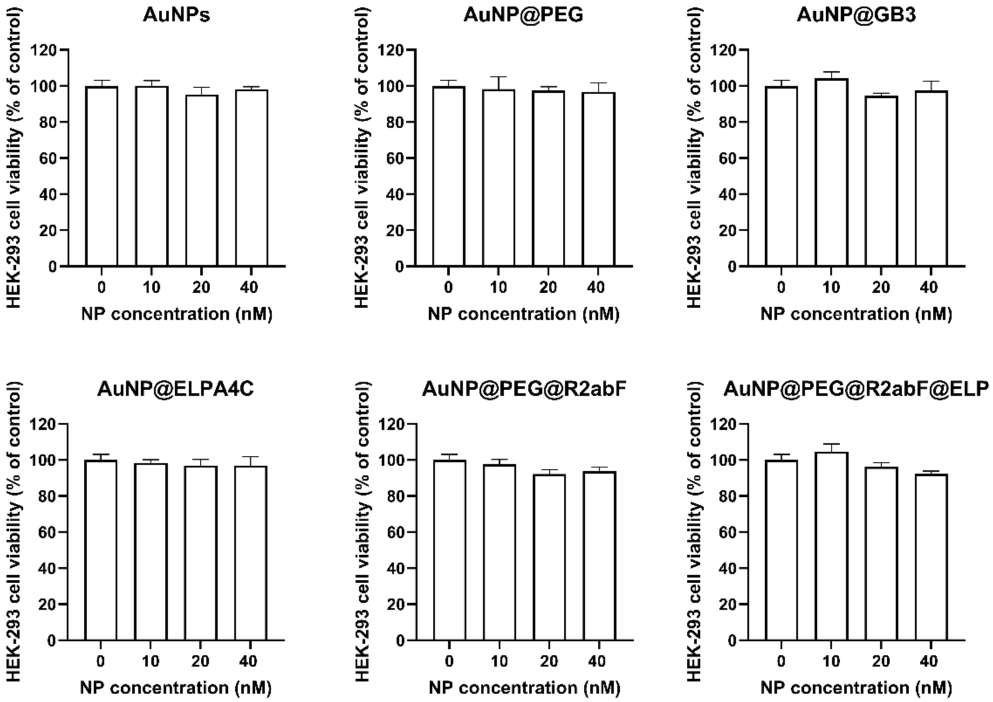

**Figure S18. Effects of different AuNP formulations on the viability of HEK-293 cells.**

The cell viability studies demonstrated that all gold nanoparticle formulations (AuNPs, AuNP@PEG, AuNP@GB3, AuNP@ELPA4C, AuNP@PEG@R2abF, and AuNP@PEG@R2abF@ELP) maintained high cell viability (>90%) in HEK-293 cells even at the highest tested concentration of 40 nM, indicating excellent biocompatibility of these nanoparticle systems. Data is expressed as the mean ± standard deviation. Treatment group changes are represented with respect to the percentage of the untreated control.
